# Evolutionary Pressures and Codon Bias in Low Complexity Regions of Plasmodia Parasites

**DOI:** 10.1101/2020.03.14.992107

**Authors:** Andrea Cappannini, Sergio Forcelloni, Andrea Giansanti

## Abstract

One of the most debated topics in Evolutionary Biology concerns Low Complexity Regions of P. falciparum, the causative agent of the most virulent and deadly form of human malaria. In this work, we analysed the proteome of 22 plasmodium species including P. falciparum. SEG predicts that proteins containing Low Complexity Regions turn out to be longer than those which are predicted to be completely complex (without Low Complexity Regions). Moreover, using some well-known bioinformatics tools such as the Effective Number of Codons, the Pr2 and a new index that we have called SPI, we have noticed how proteins that embed Low Complexity Regions are under lower selective pressure than those that do not present this type of locus. By applying the Relative Synonymous Codon Usage and other tools developed ad hoc for this study, we note, instead, how the Low Complexity Regions appear to have a non-neutral codon bias with respect to the host proteins.

## Introduction

Due to the critical implications for Human Health, the study of Single Nucleotide Polymorphisms (SNPs) and Copy Number Variations (CNVs) is a fundamental research field (Zhang et al. 2009). Tandem repeat regions (TTRs) represent a third type of genetic variation (Gemayel et al. 2012), whereby proteins host regions of reduced complexity and biased amino acid composition, also called low complexity regions (LCR) (Toll-Riera et al. 2012). Despite the diversification of the various domains of life, this type of *loci* is ubiquitously present (Kumari et al. 2015) existing in Bacteria, Archaea and Eukarya (Wootton & Federhen, 1996). Taken as a whole, the literature indicates two main mechanisms that are most likely to explain the presence of TRRs and, consequently, their propensity to undergo *indels,* whereby to cycles of expansion and contraction. The first is Replication Slippage (RS) (Guy-Franck et al. 2008; Mosbach et al. 2019; Ellegren, 2004; Gemayel et al. 2012; Saitou, 2018). RS is a mutation process that occurs during DNA replication. It involves denaturation and displacement of DNA strands with the consequent decoupling of complementary bases (Levinson & Gutman, 1987). In a nutshell, the loop out for the template strand implies a contraction whilst the same for the nascent strand implies an expansion. (Gemayel et al. 2012). The other mechanisms are recombination-like events (Gemayel et al. 2012; Ellegren, 2004; Verstrepen et al. 2005) such as unequal crossing-over and gene conversion, where some propose it as a predominant mechanism in minisatellite regions (Guy-Frank & Paques, 2000) with the predominance of replication slippage in microsatellite regions (Kokoska et al. 1998). Guy-Franck and colleagues (2008) extensively surveyed the current landscape of definitions of TRRs based on the length of the repeating unit, highlighting a lack of consensus in the definition of microsatellite where, some propose to consider a repetitive unit length of up to 5 or 10 nt either (Guy-Frank et al. 2008; Gemayel et al.2012). These *loci* have high mutational rate if compared to other DNA regions (Brinkmann et al. 1998) ranging between 10^−3^and 10^−6^per cell generation (Gemayel et al. 2012). Past research has identified some factors that would contribute to the instability of these *loci*. Gemayel and colleagues (2012) have redacted an extensive examination referring to different papers that have dealt with this topic: Legendre et al. (2007) highlighted how the presence of multiple repetitive units is the main characteristic that contributes to the instability of these regions, followed by the length and the nucleotide purity of the tract, *i.e.* a coherent succession of nucleotides not interrupted by any nitrogenous base without counterparts in the repetitive sequence (Saitou, 2018). Wanting to consider a homopolymer sequence (same amino acid) Verstrepen and colleagues (2005) have instead highlighted how the use of more synonymous codons drastically increases the stability of these *loci*. The nucleotide content also affects the instability of these regions where poly A or poly T tracts have been observed to be more stable than the corresponding poly G or poly C tracts (Gragg et al. 2002). For a more detailed examination we refer to the original work (Gemayel et al. 2012). Low Complexity Regions are thought to be the result of tri-nucleotide slippage (Levinson & Gutman, 1987; Mularoni et al. 2006). In proteins, the phenotype associated with these regions is often contradictory, whereby LCRs often have pathogenic implications, such as in neurodegenerative (Gatchel & Zoghbi, 2005) or developmental (Brown & Brown, 2004) pathologies. Nevertheless, LCRs are important for protein fitness. Shen et al. (2004) showed how the Arginine and Serine rich binding sites in the Exonic Splicing Enhancer (ESE) contribute to the assembly of pre-spliceosomes, supporting splicing and related activities. Similarly, the group of Salichs and colleagues (2009) noted that histidine-rich sites are pivotal for sub-cellular localization. More generally, incremental evidence highlights how these regions are preferentially inserted only in certain functional protein classes (Karlin et al. 2001; Alba & Guigo 2004; Faux et al. 2005) and how they do not sever their functional domains (Newfeld et al. 1994). P. falciparum is a protozoan that belongs to the *Phylum of Apicomplexa*. It represents the etiological agent of the most severe and lethal form of Human Malaria (Pizzi & Frontali, 2001). This parasite offered a fruitful source of genetic investigations, also due to its particularly high content of LCRs, mostly characterized by asparagine (N) residues. The functional role of these stretches is not fully understood yet. However, several hypotheses have been advanced. Among others, Pizzi E. and Frontali C. (2001) proposed them to be subjected, to continuous cycles of expansion and *de novo* generation representing a resource of antigenic variation ultimately leading the parasite to evade the host immune response. Such a hypothesis has been sustained by other investigators in the field (Karlin et al. 2001; Ferreira et al. 2003; Cortès et al. 2005). On the one hand and in line with a general requirement for neo-functionalization, it was proposed that long N-repetitive stretches influence the local rate of translation, thus triggering ribosome pausing and ultimately behaving as tRNA sponges that assist co-translational folding (Frugier et al. 2009; Filisetti et al. 2013). On the other hand, Forsdyke D.R. (Forsdyke, 2016) discusses the results obtained with Xue (2003), whereby similar N-repetitive stretches might play a double role at DNA level. Indeed, since the nucleotide content of P.falciparum’s genome is skewed towards AT, the N-repetitive stretches are supposed to stabilize mRNAs and prevent the corresponding genes to undergo deleterious mutations. Undoubtedly, all of the cited works have provided the scientific community with important but also conflicting explanations for the presence of LCRs in P.falciparum. Recent advances have shown how asparagine promotes the formation of amyloid structures following thermal shocks (Halfmann et al. 2011). As already explained by Muralidharan & Goldberg (2013), P. falciparum copes with extremely variable temperatures during its life cycle, *e.g*., passing from the relatively low, ambient temperature of the mosquito to the human body at about 37° C and vice versa. Moreover, Malaria is characterized by numerous cycles of fever, eventually causing the host temperature to reach > 40° C. Therefore, the continuous thermal fluctuations can lead proteins to un-as well as mis-fold that could ultimately lead to the death of the parasite. However, although Q has biochemical characteristics similar to N (Strachan et al., 2020) and is similarly prone to form prions, N reduces proteo-toxicity (Halfman et al. 2011). While showing that Heat Shock Proteins prevent the formation of aggregates and prion-like fibrils, Muralidharan et al. (2012) and Muralidharan and Goldberg (2012) proposed, among other remarkable hypotheses, a mechanism to explain how these regions can contribute to the formation of new folds and functions. In the present work, we studied the proteomes of 22 *Plasmodia* species operating on two fronts: on the one hand, the study of the Codon Usage Bias (CUB) of Low Complexity Regions by means of the Relative Synonymous Codon Usage (RSCU, Sharp & Li, 1987) and the application of Shannon’s Entropy (Shannon, 1948) provided us remarkable insights into the selective pressure acting on these regions. On the other hand, based on the presence of at least one Low Complexity Region, we stratified the proteins into LCPs (Low Complexity Containing Proteins) and not LCPs (hereafter referred to as nLCPs), the latter characterized by the absence of LCRs. Although this sub-setting was obtained by SEG predictions (Wootton & Federhen, 1996) performed using default parameters, and the results could be improved by further parameter refinement (Batistuzzi et al. 2016), our downstream results displayed internal consistency. The study of protein lengths has highlighted how, in organisms weaving a causative relationship between the two structural characteristics, LCPs are intrinsically longer than nLCPs. To address the issue of Darwinian Selective Pressures, we selected two common bioinformatics tools, i.e. The Effective Number Of Codons (Wright, 1996), using the improved version of Sun and colleagues (2012), and a Pr2 analysis (Sueoka, 1995; Sueoka & Kawanishi, 1999). In addition to these two indices, we have devised a completely new one, hereafter referred to as SPI (Selective Pressure Index), that allows to compare distances from Wright’s Theoretical Curve. The set of these three tools allowed us to highlight how, compared to nLCPs, LCPs are characterized by a more heterogeneous CUB and are further distinguished from nLCPs by a lower Selective Pressure. Our results apply consistently to any of the proteomes analysed and providing new insights about Low Complexity Regions.

## Methods

### Data Sources and general methodologies

Plasmodium proteomes from the NCBI GenBank (ftp://ftp.ncbi.nih.gov). Brazilian strain of P.vivax was considered. P.praefalciparum, P.alderi and P.o.curtisi were retrieved from PlasmoDB (https://plasmodb.org). We considered only CDSs that start with methionine (ATG) ending with a stop codon (TAG, TAA or TGA) and having a multiple length of three.

Each CDS was translated in the corresponding amino acid sequence by *nt2aa* MATLAB function. Remaining scripts were built by custom and built in MATLAB scripts. For some analyses we relied on boxplot visualization strategy. Taking into consideration two boxplots, MATLAB notch function indicates that when two notches do not overlap, medians of boxes are significantly different at the 5% significance level. (Mathworks).

We identified LCRs by SEG algorithm, triggered with standard parameters (window = 15, K1= 1.5 and K2 = 1.8) that ensures identified sequences to correspond to strongly biased sequences while, at the same time, allowing for substantial sequence diversity (Trilla & Albà, 2012).

nLCPs and LCPs often occur in groups with different numbers (see SM Supplementary Materials). Where distributions do not considerably diverge from a normal distribution, we relied on *welch-t test* (Ruxton, 2007; Derrick et al. 2016), otherwise *Mann Whitney test* was used. For multi-comparison tests we relied on *welch Anova* followed by *Bonferroni correction* (MT1) when distributions do not deviate considerably from normality, otherwise *Kruskal-Wallis test* followed by *Bonferroni correction* (MT2).

### Tandem Repeat Regions

To extract the DNA of the low complexity regions we used a customized MATLAB script in order to identify the LCRs on the proteins and extract the related DNA on the corresponding gene.

### GC, ENC, ENC plot and Pr2 analysis

We used the same rationale proposed by Forcelloni and Giansanti (2020) for the implementation of the ENC (Wright,1993) using the improved implementation of Sun et al. (2012), ENC plot, Pr2 plots (Sueoka,1995; Sueoka & Kawanishi,1999) and for the calculation of the GC content. Therefore, we refer to Forcelloni & Giansanti (2020) for further details concerning these tools. In this work, ENC and Pr2 scores are separately calculated for LCPs and nLCPs for each parasite.

### RSCU

An RSCU analysis (Sharp & Li, 1987) was carried out. RSCU measures codon usage for each codon in each codon family (Xia, 2007). Basically, RSCU is a normalized codon frequency that is expected to be 1 where there is no codon usage bias, greater that 1 when that codon is overused, minor than 1 when that codon in underused. RSCU is formally defined as follows:

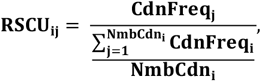

where *CdnFreq_j_* refers to a codon frequency *j*; *NmbCdn_i_* is the degeneracy of the codon family *i*; 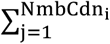 CdnFreq_i_ is the sum of the occurrences of the codons in that codon family that encode for the same amino acid. Worth noting, the maximum value that can be expected from a codon depends on the codon family to which it belongs. Therefore, if the degeneracy of a codon family is n then the maximum expectation will be n. We introduced a modification to the original version of RSCU proposed by Sharp & Li (1987), in line with the division of the 6-fold codon families into 4-fold and 2-fold proposed by Sun and colleagues (2013) in their improved version of ENC that we use in our ENC analysis. We neglect on purpose ATG (M) and TGG (W) since being expressed by only one codon there cannot be bias. We computed, for each species, RSCU analysis on the overall codon content of Low Complexity Regions.

### Shannon Entropy

To compare the complexity of Tandem Repeat Regions we thus relied on *Shannon’ Entropy* (Shannon,1948) computed as follows:

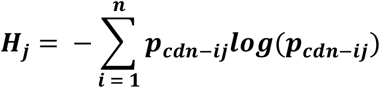

where 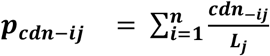 that is the sum of the *i-th* codon in the *j-th* tandem repeat of length *L_j_* Entropy is used to represent in a compact way Tandem Repeat Regions and their bias towards a few or more codons.

### SPI

The Effective Number of Codons describes the codon bias of a gene. Moving away from the curve, the distance from the Wright’s Theoretical Curve provides an estimate of the extent to which Mutational Bias and Selective Pressure affect the codon bias of that gene (Novembre, 2002). On the one hand, the same ENC score be associated with different distances from the curve, and the other, the same distance in absolute value, as the GC3 varies, can represent a diverse deviation from the expected value given by Wright’s Theoretical Curve. The distances in absolute value are not equivalent. Therefore, we propose the SPI (Selective Pressure Index) defined as follows:

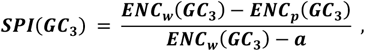

where *ENC_w_*(*GC*_3_) is the value of the Wright’s theoretical curve in correspondence of a *GC*_3_ *ENC_p_*(*GC*_3_) is the ENC value for the gene in correspondence of that *GC*_3_ value and a = 20. SPI weights the shift of the coding sequence from the null model of no-codon preference (the situation where a gene lies on the curve) with the situation of extreme bias, namely where just one codon is used for each amino acid, for that *GC*_3_. When a gene lies on the theoretical curve SPI is expected to be 0 (no distance from the WTC). Otherwise, when a gene is extremely subject to selection SPI is expected to be 1 (situation of extreme bias).

### Cladistic and Organism Overview

We briefly describe how the parasites are phylogenetically related and why we placed them in a subgroup rather than in another. Let us point that extending the knowledge about the various radiations that brought to the occurrence of these subgroups is beyond our scope. P.falciparum has been placed together with the other parasites belonging to the *monophyletic subgenus* termed *Laverania* (Otto et al. 2018;Liu et al. 2016) that comprehends P.gaboni and P.reichnowi (that infect chimpanzees), P.praefalciparum and P.alderi (that infect gorillas). Noteworthy, P.falciparum is the only that successfully infects humans (Otto et al. 2018) but despite it was for a long time considered a human specific pathogen it also infect gorillas, rising concerns about possible reciprocal host transfer (Prugnolle et al. 2010). As far as the Asian monkey parasites are concerned, we retrieved data for P. vivax that represents a serious threat for human health given its extremely wide geographical coverage (Howes et al. 2016). P. knowlesi was recently recognized as a human pathogen (Rich & Xu, 2011) given a major outbreak of human infection in 2004 (Singh et al. 2013). Until this focus, human contagions were thought rare (Faust & Dobson, 2015). Moreover, we considered P.cynomolgi and P.coatney (Galinski & Barnwell, 2012), P.inui (Galland, 2000) and P.gonderi (Arisue et al. 2019). We refer to these parasites as S*imian Plasmodia*. To have a blueprint about evolutionary and quantitative diversification of LCRs we considered also murine rodents plasmodia (*Vinckeia Subgenus*). We retrieved P.vinckei (Carter & Wallinker, 1975), P.petteri, P.chabaudi, P.berghei and P.yoelii (Garnham 1964). Evidence addresses the evolutionary origin of P. falciparum with avian plasmodia (Waters et al. 1991; Waters et al. 1993 a–b; McCutchan et al. 1996; Escalante & Ayala 1994). To extend our efforts we considered the two available specimens of the Haemamoeba Subgenus (Corradetti et al. 1963): P. gallinaceum, proposed as a possible ancestor of P.falciparum (Waters et al. 1991;Escalante et al. 1998), and P.relictum that is one of the most geographically widespread malaria parasites for birds (Valkiunas, 2005). We refer to these parasites as *Haemamoeba Subgenus.* Lastly, we considered P.ovale wallikeri (P.ovale), the etiological agent of tertian malaria, (Collins & Jeffery, 2005), P.ovale curtisi (P.o.curtisi) (Kristan et al. 2019) and P.malariae causing quartan malaria (Collins & Jeffery, 2007). We refer to these parasites as Human Infectious Plasmodia (*HIPs*).

## Results

### GC – Content Influence

Having the possibility to make a statistical comparison among several parasites, we reflected on the role of Guanine and Cytosine content, in order to understand the degree to which LCRs of P.falciparum, and more generally of the other parasites, are influenced by genomic biases. We considered the Laverania Plasmodia as a yardstick for our analysis. In **Tab.1** we summarize the statistics from various experiments we performed during the analysis. The first column collects the *Pearson* correlation coefficients between the amount of Low Complexity Regions contained in proteins and their length. Next, the average Guanine and Cytosine content of each protozoan and the quantity of low-complexity regions possessed by each of them. All the correlations are significant (p < 0.01). We have divided the statistics of each column by parasite group. Thus, each column was divided into 5 distributions. MT2 applied to the three distributions, indicates that Laverania, Haemamoeba and Vinckeia subgroups do not differ significantly in Guanine and Cytosine content. However, the Vinckeia have significantly lower correlations than the P. falciparum’s group. We report the same also as regards the contents in Low Complexity residues. Although the Haemamoeba and Laverania Plasmodia do not differ significantly in correlation coefficients, the statistical pauperism of the group could lead to misleading results. In a nutshell, genomic bias does not seem to play a particular selective advantage although we do not deny its role in the choice of synonymous codons.

**Tab.1.** In the first column, the Pearson correlation coefficients of 22 distributions are collected in which the length of the proteins and the amount of LCR present in them is correlated. The second column collects the average Guanine and Cytosine content of each parasite. The third column collects the number of low complexity residues present within each organism as identified by SEG.

| Organism | Pearson-correlation coefficient | GC content | LC content (amino – acids count) |
| --- | --- | --- | --- |
| *P.falciparum* | r = 0.70 | 0.23 | 130020 |
| *P.praefalciparum* | r = 0.71 | 0.23 | 125491 |
| *P.reichnowi* | r = 0.71 | 0.23 | 133878 |
| *P.alderi* | r = 0.67 | 0.22 | 105572 |
| *P.gaboni* | r = 0.70 | 0.22 | 102893 |
| *P.vivax* | r = 0.35 | 0.48 | 28753 |
| *P.knowlesi* | r = 0.26 | 0.40 | 19693 |
| *P.coatney* | r = 0.32 | 0.44 | 15273 |
| *P.cynomolgi* | r = 0.42 | 0.42 | 26101 |
| *P.inui* | r = 0.26 | 0.44 | 16109 |
| *P.fragile* | r = 0.22 | 0.44 | 19378 |
| *P.gonderi* | r = 0.39 | 0.26 | 31857 |
| *P.yoelii* | r = 0.33 | 0.24 | 29765 |
| *P.berghei* | r = 0.38 | 0.24 | 28753 |
| *P.petteri* | r = 0.21 | 0.27 | 11737 |
| *P.vinckei* | r = 0.16 | 0.26 | 11324 |
| *P.chabaudi* | r = 0.26 | 0.28 | 11265 |
| *P.gallinaceum* | r = 0.46 | 0.21 | 49707 |
| *P.relictum* | r = 0.37 | 0.21 | 39707 |
| *P.malariae* | r = 0.45 | 0.30 | 31876 |
| *P.ovale* | r = 0.43 | 0.36 | 43101 |
| *P.o.curtisi* | r = 0.30 | 0.32 | 32005 |

### RSCU

We wanted to study the Codon Bias of the Low Complexity Regions to derive information about the selective pressures acting on their synonymous codons. To do this, we used Relative Synonymous Codon Usage (RSCU, Sharp & Li, 1987) applied on the overall Codon Content of LCRs. We have deliberately neglected ATG (M) and TGG (W) because being encoded by only one codon they cannot have bias. The results are displayed in **Fig. 1**. Data is standardized along each column. If a codon has a value above the column mean it is represented in red. Otherwise green. Specifics regarding the clustering algorithm and distance are provided in the image caption. The heatmap indicates that *Protozoa* can be distinguished in two main lines, although there are differences within each group. The Laverania, Haemamoeba and Vinckeia Plasmodia mainly prefer codons whose wobble position is occupied by Adenine or Thymine, when compared to the remaining subgroups (HIPs and Simian Plasmodia). The latter, on the contrary, have larger RSCU values for codons whose wobble position is represented by Cytosine or Guanine, if compared to the previous species. We wanted to investigate further about the RSCU values within each species in order to notice any more specific preferences in respect of the wobble positions. To do this, we compared the RSCU values of codons ending with a purine (A or G) and codons ending with a pyrimidine (T or C). The Relative Synonymous Codon Usage is a normalized index that presents a spectrum of different maximum values that depend on the degeneracy of the codon family. Therefore, we have normalized the RSCU values with respect to the number of codons of the family they belong to (*Mann Whitney*). As expected, Laverania, Vinckeia and Haemamoeba Plasmodia follow the copycat, showing a clear preference for A over G and T over C in the wobble position (p <0.01), as it is also for HIPs (p<0.01). We find the same pressures in P.gonderi (p <0.01). Interestingly, we do not find the same stark split in many of the *Simian Plasmodia*, such as P.vivax, P.cynomolgi, P.fragile and P.inui whose wobble position appears to be contested in a tug-of-war between both purines and pyrimidines, not finding significant differences (*p>0.05*). Evolutionary pressures are slightly different in P.coatney and P.knowlesi. The first has preference for G over A and T over C (p <0.05). The second presents wobble positions better suited to its GC content by preferring G to A and C to T (p <0.05). RSCU table is provided in the Supplementary materials (*RSCU.xlsx*). Overall, we obtained information about the synonymous codons used in these organisms. However, RSCU is a normalized index which neglects the quantitative contribution of a codon family which, in contrast, could be selectively avoided.

**Fig 1.**
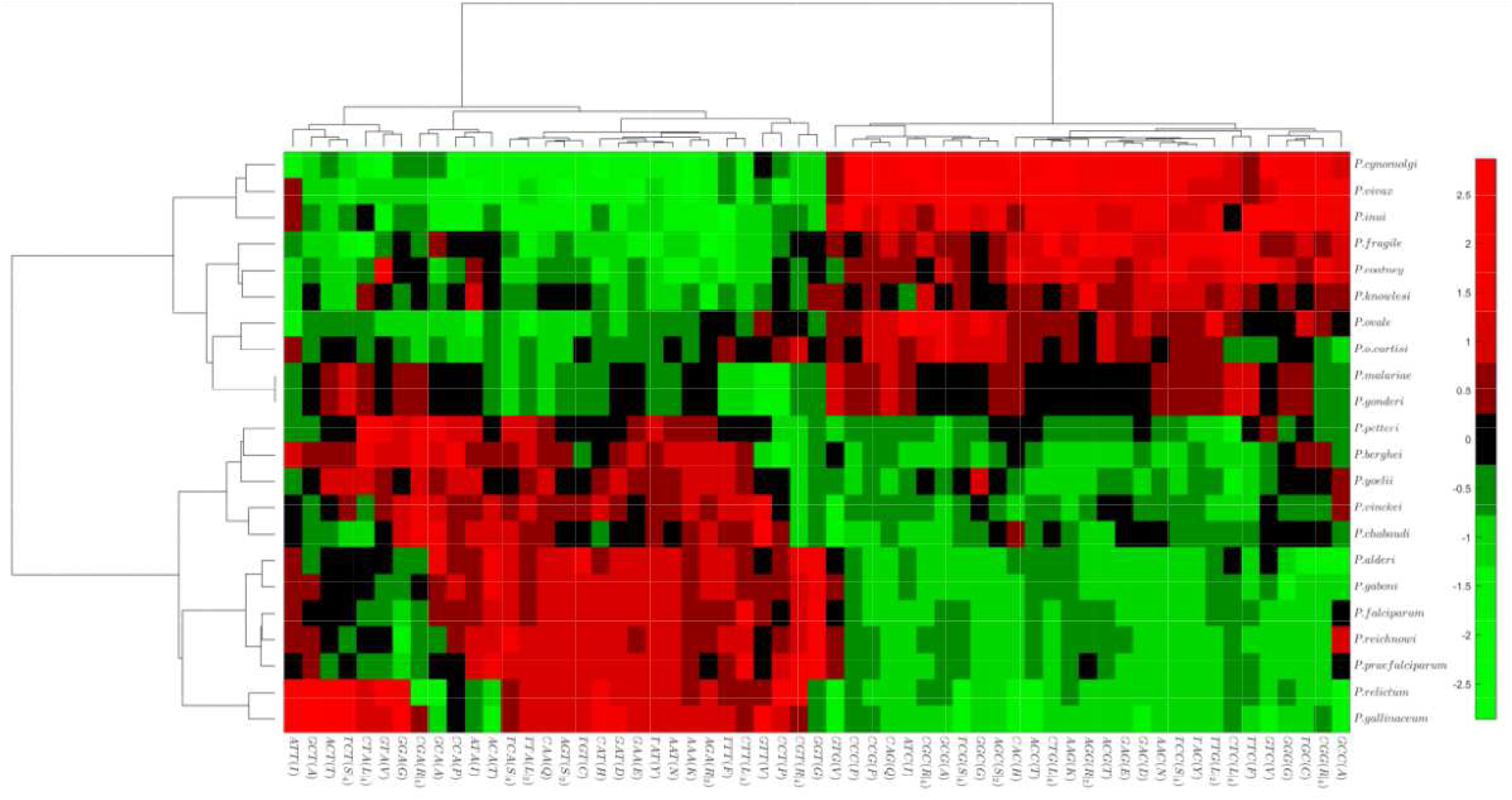
Heatmap of the RSCU values of the Low Complexity Regions. Codons are placed on the columns. Plasmodia on the rows. Data is standardized along the columns. **Ward’s Linkage** was used. Row and column pairwise distance was calculated with **Standardized Euclidean Distance**. Data were standardized along rows. The **MATLAB clustergram function** was used.

### Shannon Entropy and Complexity of Tandem Repeats Regions

We investigated the codon composition of the LCRs of each plasmodium using Shannon Entropy to understand their structural arrangement and derive a rationale for inferring the presence of different selective pressures. **Fig. 2(A)** illustrates the distributions of the 5 parasitic groups. Best fit parameters and models are left in the image caption. The elbow of the regressions shows that, on average and for the same length, the LCRs of the Laverania Plasmodia are less complex than those of the other parasitic groups, especially HIPs and Simian spp. To have a better understanding of the phenomenon we have studied the single distributions as shown in **Fig. 2 (B)**. Preliminary observation of the violin plots delineates how Plasmodium species arrange their TRR differently, which denotes, as entropy increases, a greater number of codons along the same TRR. We have applied MT1. The Bonferroni Correction indicates that the Laverania Plasmodia use fewer codons (synonyms / non synonyms) along the extension of their LCRs as opposed to other species whose Low Complexity Regions appear more complex. The only parasite for which the comparison with Laverania Plasmodia is insignificant is P.berghei. **Fig. 2 (C)** shows the correlation between the average complexity of TRRs and the average Guanine and Cytosine content of each organism. In general, average TRR complexity increases with the GC content even if the fluctuations suggest the presence of other forces in determining the nucleotide composition of these regions.

**Fig 2.**
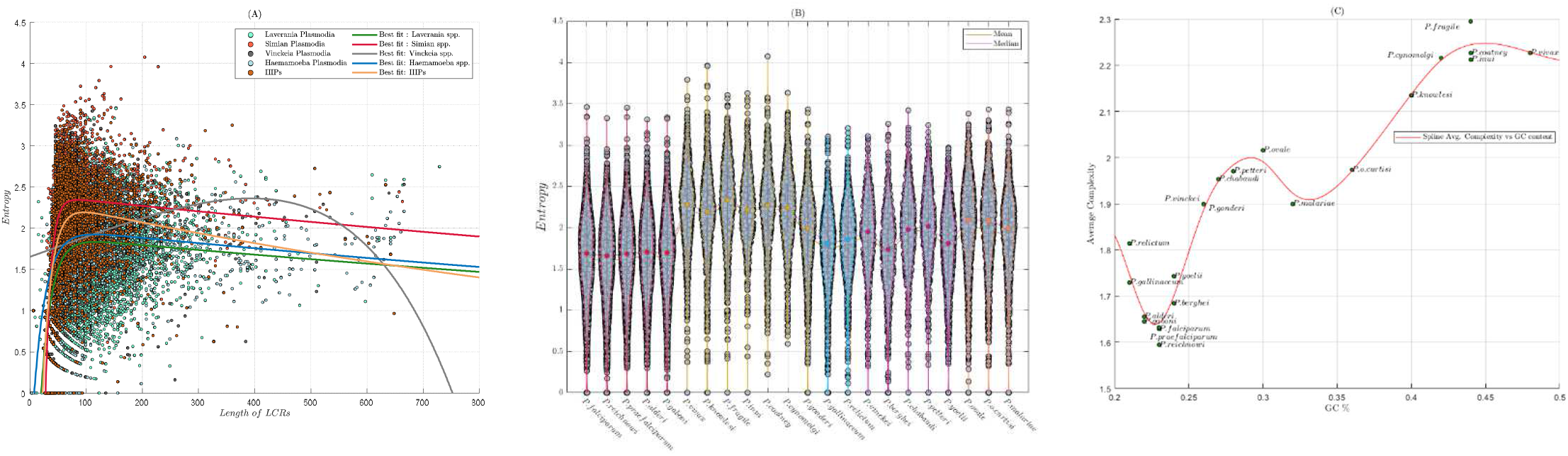
**(A)** Regression of Entropy vs LCRs length. Length of LCRs is measured in nucleotides. Model parameters are provided with 95% coefficient bounds **Laverania(x) = a*exp(b*x) + c*exp(d*x): a =** 1.918 (1.885, 1.951),**b =** −0.0003328 (−0.0004543, −0.0002113), **c =** −5.435 (−5.793, −5.078), **d =** − 0.05285 (−0.05514, −0.05056); **Simian(x) = a*exp(b*x) + c*exp(d*x): a =** 2.41 (2.368, 2.452), **b =** −0.0002985 (−0.0004451, −0.0001519), **c =** −39.41 (−47.79, −31.02), **d =** −0.1012 (−0.1077, −0.09478); **Vinckeia(x) = a*exp(b*x) + c*exp(d*x): a =** −1.006e+04(−4.838e+12, 4.838e+12), **b =** 0.002788 (−52.56, 52.57),**c =** 1.006e+04 (−4.838e+12, 4.838e+12),d = 0.002788(−52.55, 52.56); **Haemamoeba(x) = a*exp(b*x) + c*exp(d*x): a =** −2.832 (−3.196, −2.469),**b =** −0.04565 (−0.05072, −0.04057), **c =** 2.018 (1.956, 2.08), **d =** −0.0003467 (−0.0004884, −0.000205); **HIPs(x) = a*exp(b*x) + c*exp(d*x): a =** −9.49 (−10.81, −8.165), **b =** −0.06278 (−0.0674, −0.05817), **c =** 2.357 (2.291, 2.423), **d =** −0.0006497 (−0.0008514, −0.0004479) **Fig. 2(B)** Illustration of the average complexity of each Low Complexity. The mean of each distribution is represented by a data point of the same colour as the outline of the violin plot. The trend of the means and medians is represented by the yellow and red lines respectively. Violin Plot Function has been taken from <u>*GitHub-Matlab*</u> **(C)** Trend of the sample averages relative to the distributions of **Fig. 2 (B)** with respect to the average Guanine and Cytosine content of each parasite

### Codon Bias

We have analyzed from a quantitative point of view the codon bias of the Low Complexity Regions since the RSCU cannot return a quantitative information regarding the use of codons (**Fig.3**). *P_Cdn_* represents the ratio between the total quantity of a codon and the total number of codons present within the Low Complexity Regions of a parasite. All the species, although in different proportions, show an increase in the codons used less commonly than those with higher percentages, unlike the Laverania Plasmodia which have a skewed distribution towards a few species of codons.

**Fig 3.**
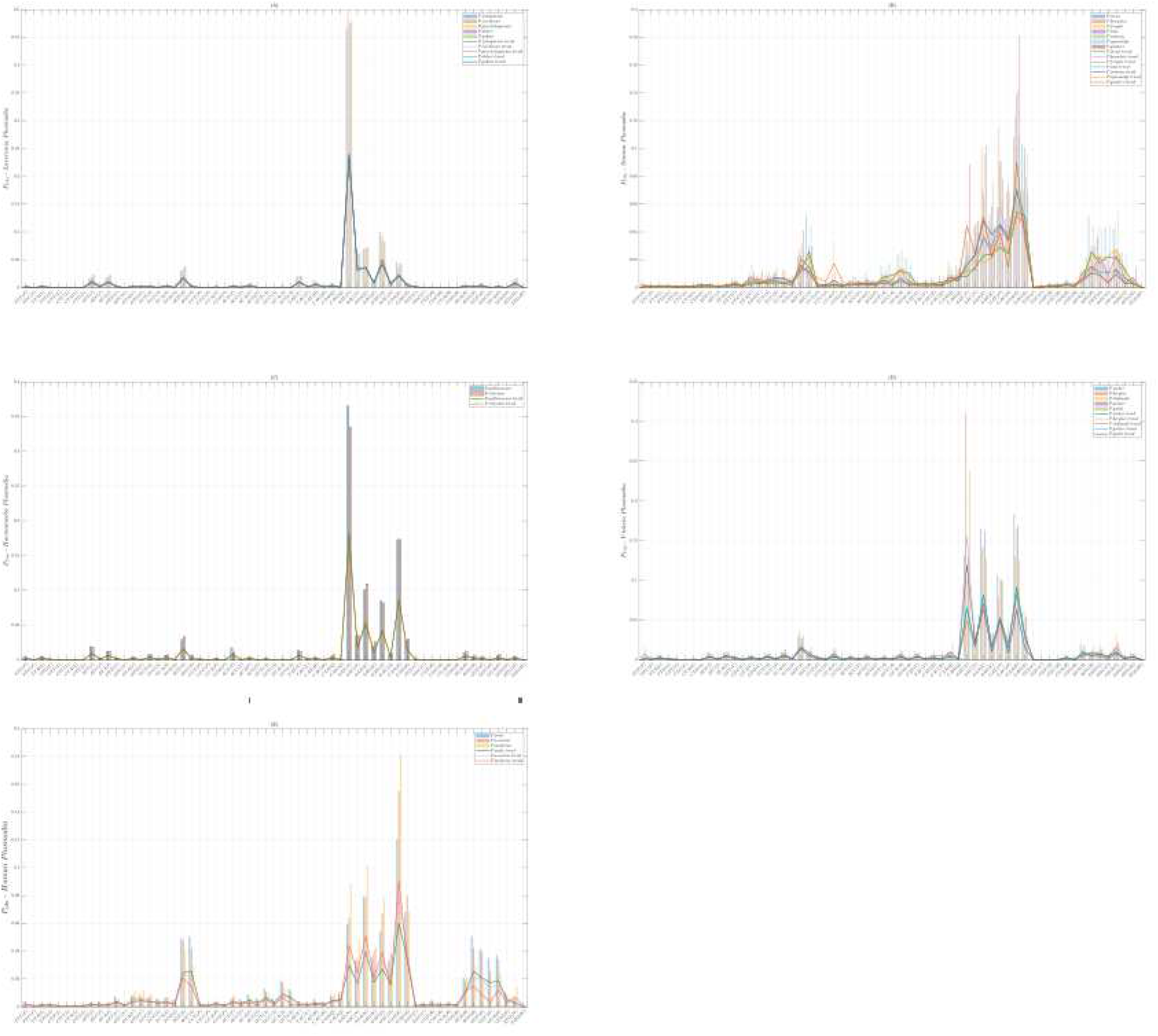
Percentage representation of the CUB in the various parasitic subgroups. The amount of each codon was divided by the total amount of residues present in the LCRs of each parasite. **(A) Laverania Subgenus (B) Simian Plasmodia (C) Haemamoeba Subgenus (D) Subgenus Vinckeia (E) HIPs.** Each peak represents half of the bar it refers to.

### Protein Length

Given the positive correlations between LCRs and protein length we observed in 1^st^ paragraph, we analyzed the protein length of each Plasmodium by stratifying proteomes into two distributions, i.e., LCPs and nLCPs. We compared the two distributions in each organism (Welch t-test) (**Fig. 4 (a)**), represented in red and green respectively. LCPs emerge to be significantly longer than nLCPs in each parasite (p << 0.01). We have reflected on the linear extension of the LCRs. Considering LCRs length (See Supplementary Materials), individually, these do not exceed 250 amino acids in length. However, it is not uncommon to find more than one Low Complexity Region within the same protein. Therefore, for the avoidance of doubt, we studied the proportion between LCRs and protein lengths to understand with certainty whether, on average, the length of LCPs was not only an artifact due to the presence of these stretches (**Fig. 4 (b)**). From what emerges, LCRs occupy a fairly small space, rarely reaching 6% of the entire linear surface of the polypeptides. We therefore deprived the LCPs of their Low Complexity Regions and repeated the experiment by comparing LCPs and nLCPs (**Fig. 4 (c)**) (Welch t-test). LCPs emerge intrinsically longer than nLCPs (p << 0.01). Even if the graph shows a lack of statistically significant difference between the length distributions of LCPs of the various species, we wanted to apply MT1 to these to understand if there could be differences such to explain the overabundance of LCRs typical of the Laverania Plasmodia. Apart from P. gonderi and P.malariae which appear to have the longest LCPs among all parasites and P.berghei which instead appears to have the shortest, no differences emerge such as to justify the overabundance of LCRs typical of the Laverania Plasmodia. Overall, LCPs emerge longer *de facto* than nLCPs which weaves, in general, a causative link between protein length and the presence of Low Complexity Regions.

**Fig 4.**
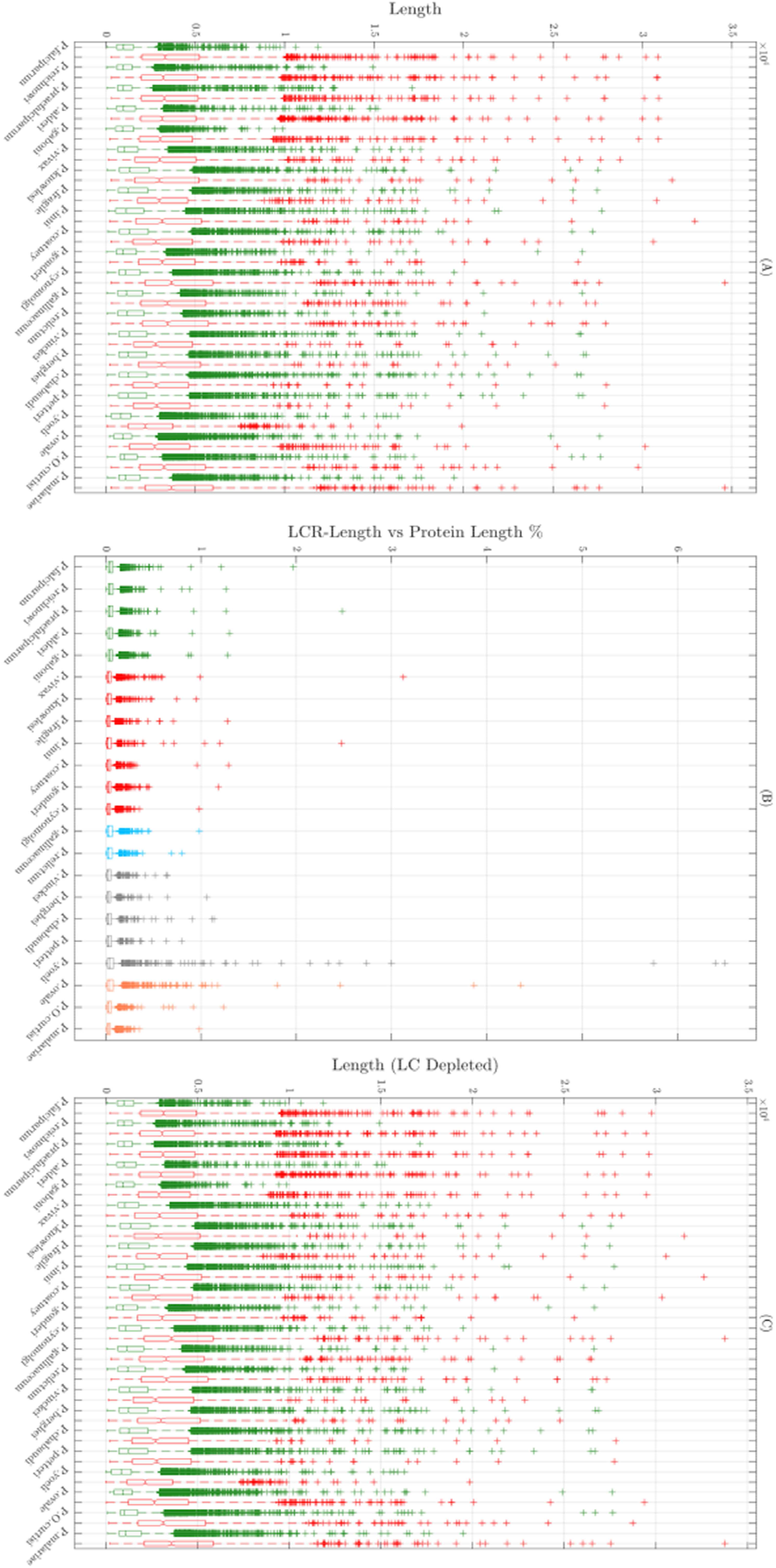
**(A)** Pairwise comparison of LCPs and nLCPs in each parasite. **(B)** Relationship between the length of the LCRs and the length of the host protein. **(C)** Pairwise comparison of LCPs, deprived of LCRs, and nLCPs. The LCPs are represented in red. The nLCPs are represented in green.

### ENC

We relied on the Effective Number of Codons (Wright,1990). We utilized the improved implementation proposed by Sun and colleagues (2013). ENC is a widely used index that does not require a reference set as the Codon Adaptation Index (CAI) does (Sharp & Li, 1987). We calculated separately ENC scores for LCPs and nLCPs. They are represented in red and green, respectively. Laverania, Vinckeia and Haemamoeba Subgenera (**Fig.5 (A) (C) (D)**) show off similar distributions placed on the left side of the ENC plane, highlighting a low GC content in Wobble codon position. Interestingly, nLCPs of P.yoelii, despite mainly grouping on the left side of the ENC plot, distribute throughout the plane following the Wright’s Theoretical Curve. HIPs (**Fig.5(B)**) have a *GC*_3_ content halfway between former Subgenera and Simian Plasmodia whose ENC plots, in turn, are in line with other works (Gajbhiye et al. 2017; Yadav & Swati, 2012). Remarkably, P.vivax and P.cynomolgi have a portion of their nLCPs positioned on the left side of the ENC plane (**Fig.6)**, that further emphasises their close phylogenetic relationship (Sanger Institute). All but Simian Plasmodia, show ENC values that distribute linearly with *GC*_3_. We collected the correlation coefficients (Pearson) of ENC vs *GC*_3_ in **Tab. 2** together with regression slopes of both nLCPs and LCPs. Given the shape of its distributions we added also P.gonderi. All the correlations are significant (p<0.01). We compared slope distributions of nLCPs and LCPs (Mann Whitney). LCPs regressions are on average represented by steeper slopes (p<0.01). Mann Whitney test indicates a greater influence of *GC*_3_ pressure on LCPs. In **Tab. 3** we collected the *r*^2^ of regression curves we calculated for Simian Plasmodia. Models and parameters, calculated with 95% confidence bounds, are provided in the caption of **Fig.6**. Generally, the regression curves of LCPs tend to overlook those of nLCPs. To add statistics to our analysis, we compared the ENC scores of LCPs and nLCPs parasite-wise (**Fig. 7**) (Welch t-test). Most of the comparisons show that the LCPs have a more redundant codon bias compared to nLCPs (p<<0.01). The test fails only for P.chabaudi, P.yoelii and P.berghei (p>0.05) even if their medians differ with 95% confidence. In a nutshell, LCPs have a more redundant CUB which suggests a stronger action of mutational bias than Darwinian Selective Pressure (positive / negative).

**Fig 5.**
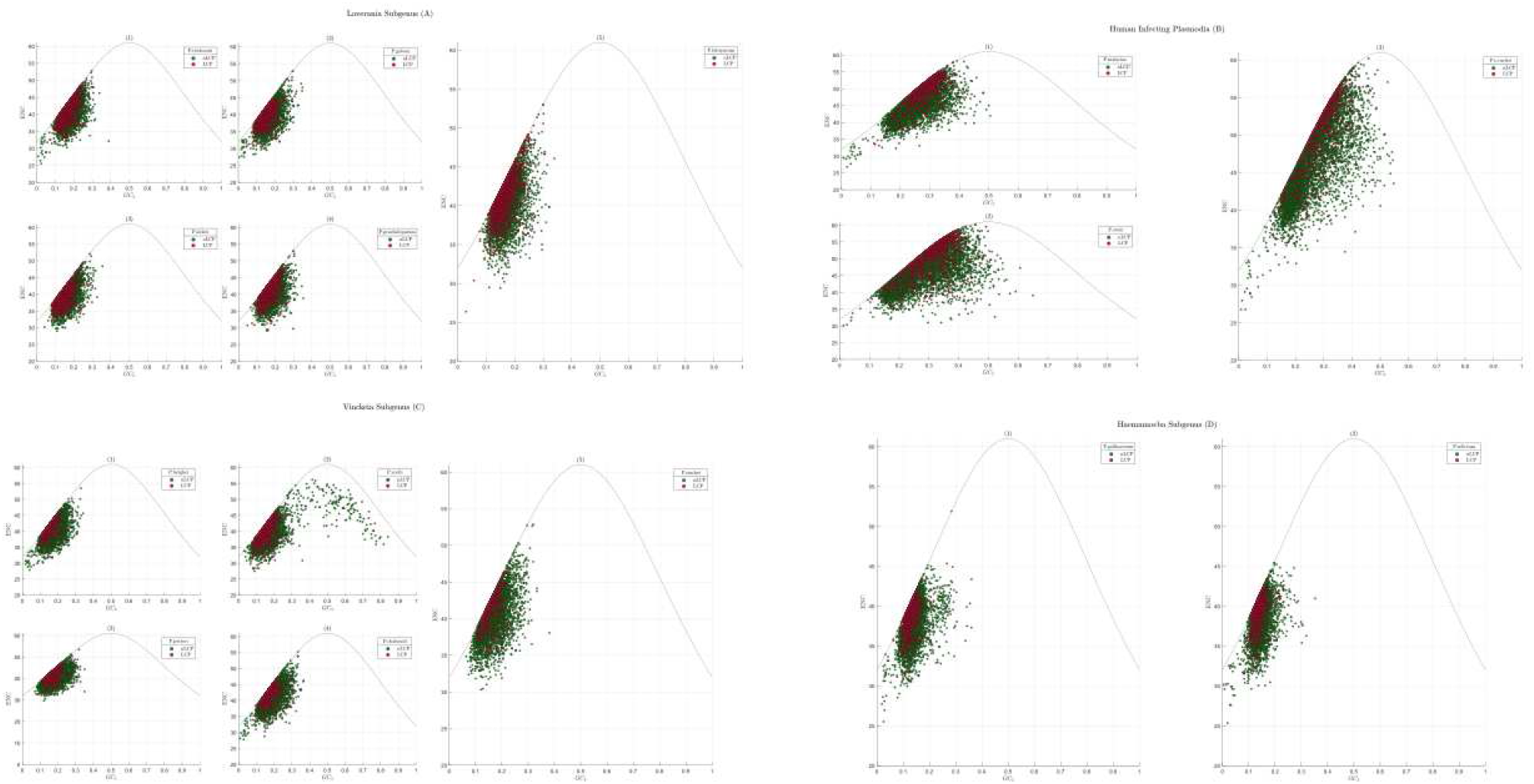
Illustration of ENC plot for **Laverania Subgenus: (1)** P.reichnowi **(2)** P.gaboni **(3)** P.alderi **(4)** P.praefalciparum **(5)** P.falciparum; **Human Infectious Plasmodia: (1)** P.malariae **(2)** P.ovale **(3)** P.o.curtisi; **Vinckeia Subgenus: (1)** P.berghei**(2)** P.yoelii **(3)** P.petteri **(4)** P.chabaudi **(5)** P.vinckei; **Haemamoeba Subgenus: (1)** P.gallinaceum **(2)** P.relictum. Globally ENC distributions cluster in the left region of the ENC plane. LCPs are represented in Red. nLCPs are represented in green.

**Fig 6.**
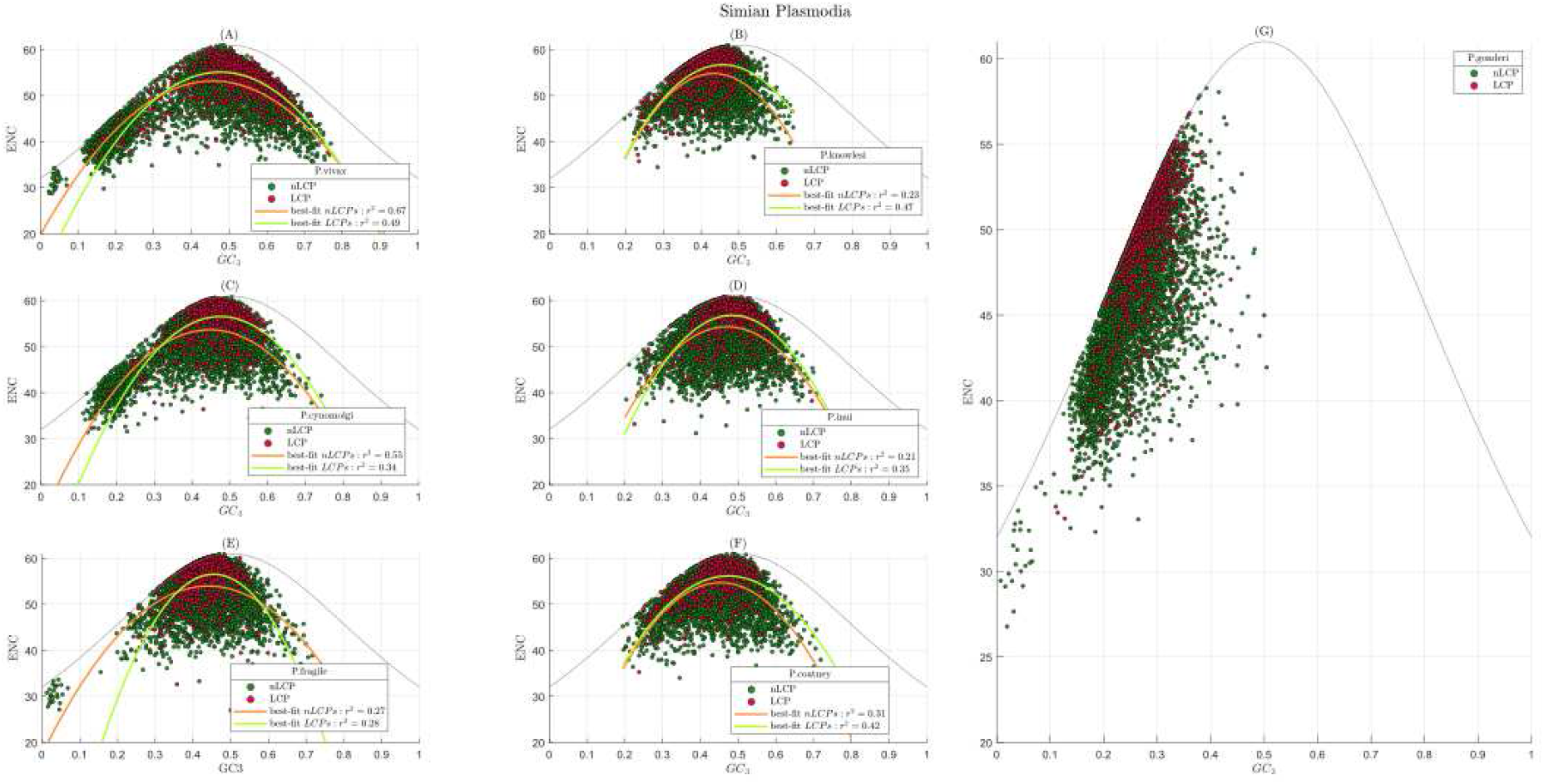
Illustration of the ENC Plots of **Simian Plasmodia**. We used a f(x) = p1*x^2 + p2*x + p3 model. Best Fit’s parameters **(BFPs)** are provided with the 95% confidence bounds. In red LCPs, in green nLCPs. In orange the best fit for nLCPs. In light green the best fit for LCPs. **(A) P. vivax: nLCPs’ BFPs: p1** = −160.3 (−164.3, −156.3), **p2** = 146.5 (143.3, 149.7),**p3** = 19.7 (19.13, 20.27);LCPs’ BFPs: **p1** = −197.6 (−211.5, −183.7),**p2** = 188.5 (175.5, 201.4),**p3** = 10.18 (7.227, 13.14) **(B) P.knowlesi: nLCPs’ BFPs: p1 =** −334.7 (−352.9, −316.5),**p2 =** 289.9 (274.4, 305.4),**p3 =** −7.969 (−11.24, −4.701); **LCPs’ BFPs: p1 =** −298.4 (−351, −245.8),**p2 =** 272.7 (232.2, 313.2), **p3 =** −5.609 (−13.32, 2.101) **(C) P.cynomolgi: nLCPs’ BFPs: p1 =** −207 (−214.2, −199.8),**p2 =** 185.1 (179.6, 190.7),**p3 =** 12.28 (11.23, 13.32); **LCPs’ BFPs: p1 =** −259 (−291.5, −226.5),**p2 =** 245.9 (217.5, 274.3),**p3 =** −1.82 (−8.019, 4.379) **(D) P.inui: nLCPs’ BFPs: p1 =** −268.7 (−284.1, −253.3),**p2 =** 253.2 (239.1, 267.3),**p3 =** −5.331 (−8.551, −2.111);LCPs’ BFPs: **p1 =** −320.5 (−363.3, −277.8),**p2 =** 310.3 (271.1, 349.6),**p3 =** −18.31 (−27.29, −9.327) **(E) P.fragile: nLCPs BFPs: p1 = −188.2 (−197.4, −179),p2 = 165.9 (158.1, 173.6),p3 = 17.42 (15.75, 19.09);LCPs’ BFPs: p1 =** −416.8 (−476.9, −356.6)**,p2 =** 379.3 (327.3, 431.2)**,p3 =** −29.75 (−40.93, −18.57); **(F) P.coatney: nLCPs’ BFPs: p1 =** −280.9 (−293.3, −268.6)**,p2 =** 254.8 (244, 265.6)**,p3 = −**3.065 (−5.406, −0.724);**LCPs’ BFPs**: **p1 =** −246.7 (−289.4, −203.9), **p2 =** 234 (198.5, 269.5),**p3 =** 0.6524 (−6.597, 7.902) **(G)** P.gonderi

**Tab.2.** Correlation coefficients between ***GC*_3_** content of each protein and their ENC scores.

| Organism | nLCP | LCP | nLCPs-slope | LCPs-slope |
| --- | --- | --- | --- | --- |
| *P.falciparum* | r = 0.56 | r = 0.65 | 43.9 | 55.4 |
| *P.praefalciparum* | r = 0.57 | r = 0.62 | 46.8 | 51.5 |
| *P.reichnowi* | r = 0.62 | r = 0.62 | 49.2 | 52.9 |
| *P.alderi* | r = 0.61 | r = 0.60 | 47.4 | 48.1 |
| *P.gaboni* | r = 0.59 | r = 0.53 | 44.2 | 41.7 |
| *P.yoeli* | r = 0.54 | r = 0.52 | 23.4 | 44.5 |
| *P.berghei* | r = 0.57 | r = 0.62 | 43.3 | 51.5 |
| *P.petteri* | r = 0.54 | r = 0.47 | 41.4 | 41.5 |
| *P.vinckei* | r = 0.57 | r = 0.54 | 41.9 | 43.8 |
| *P.chabaudi* | r = 0.60 | r = 0.51 | 43.9 | 39.3 |
| *P.gallinaceum* | r = 0.42 | r = 0.43 | 30.8 | 37.8 |
| *P.relictum* | r = 0.41 | r = 0.41 | 37.7 | 44.5 |
| *P.malariae* | r = 0.68 | r = 0.72 | 50.4 | 63.5 |

**Tab.3.** ENC vs *GC*_3_ *r*^2^ values. Simian Plasmodia

| Organism | nLCP | LCP |
| --- | --- | --- |
| **P.vivax** | $r^2 = 0.67$ | $r^2 = 0.50$ |
| **P.knowlesi** | $r^2 = 0.23$ | $r^2 = 0.47$ |
| **P.inui** | $r^2 = 0.20$ | $r^2 = 0.35$ |
| **P.cynomolgi** | $r^2 = 0.56$ | $r^2 = 0.34$ |
| **P.coatney** | $r^2 = 0.32$ | $r^2 = 0.42$ |

**Fig 7.**
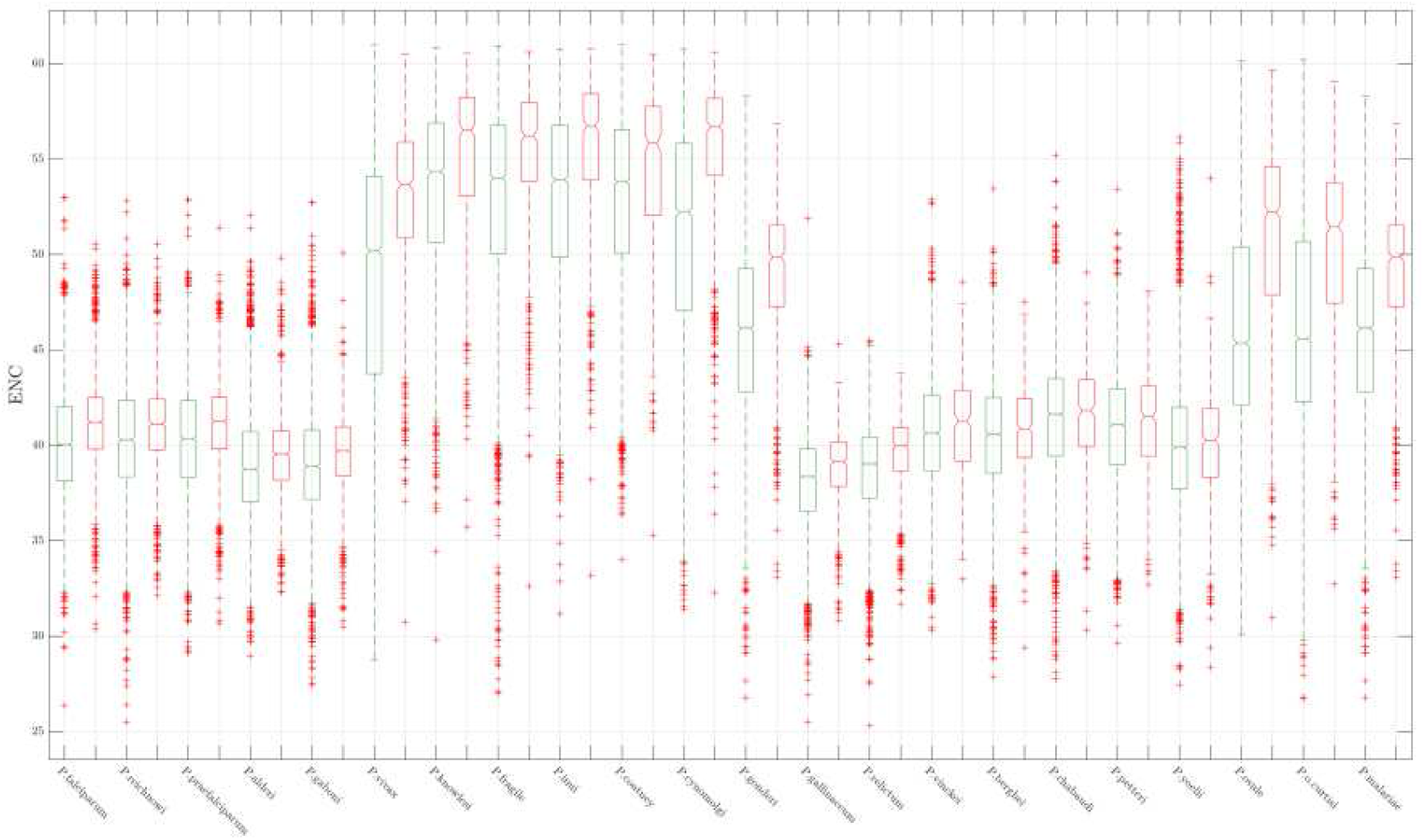
Pairwise **(Welch t-test)** comparison of the ENC distributions of LCPs and nLCPs in each parasite. Consistent with what was done in the previous graphs, the LCPs are represented in red. Similarly, nLCPs are represented in green. The medians of each boxplot pair differ with 95% statistical significance. (<u>Mathworks</u>).

### SPI

Analysis of the Effective Number of Codons has revealed some very interesting details related to the codon bias of LCPs and nLCPs which, on average, seem to differ in the extent to which selective pressure and mutational bias affect the choice of their codons. The problem of the ENC is that the distances in absolute value from the Wright Theoretical Curve are not equivalent to each other and therefore two equal ENC values can represent two different contributions of the selective pressure and of the mutational bias. In this regard we have applied the SPI (**Fig.8**) repeating what we have already done in **Fig.7** (Welch t-test). The SPI analysis corroborates what was primarily observed by ENC. Indeed, LCPs, on average, emerge to be under a lower selective pressure than nLCPs (p<<0.01) diverging less from the Wright Theoretical Curve in each parasite. Medians of each pair differ with 95% confidence. We evaluated the hypothesis that a lower selective pressure could favour a greater pervasiveness of LCRs in the Laverania Subgenus. We therefore applied MT1 on LCP distributions. The test returns contradictory values. In fact, although the Simian group shows the highest SPI values compared to all the other parasites considered here, some parasites of the Vinckeia and Haemamoeba groups show values similar to or lower than the P. falciparum group. Thus, seeing that LCPs of AT-rich groups undergo similar patterns of selective pressure, the hypothesis that greater mutational bias increases LCRs lapses. For a better understanding we refer to the interactive MATLAB plot left in the Supplementary Materials. Duret and Mouchiroud (1999) noted a negative correlation between protein length and codon bias in *C.elegans*, *D.melanogaster*, and *A.thaliana*. We therefore decided to retrace what they did using the SPI (**Fig. 9**). In the proposed plan, each data point (Laverania Plasmodia) represents a protein where the length (abscissa) is measured in nucleotides. SPI values are placed on the ordinate. Regression model (fit MATLAB function) and parameters calculated with 95% confidence interval are provided in the image caption. Again, we point out that the LCPs have been deprived of their LCRs. The graph highlights how the contribution of the mutational bias tends to increase with the length of the protein for both nLCPs and LCPs. Selective Pressure decreases as protein length increases. Similar trends are also reported for the other parasitic groups whose graphs and models are provided in Supplementary Materials. Recalling the correlations between LCRs and protein length observed in the first paragraph, these trends contain another hidden information, namely that the lower the selective pressure, the more LCRs are inserted. This weaves, more clearly, a relationship between Mutational bias and Low Complexity Regions.

**Fig 8.**
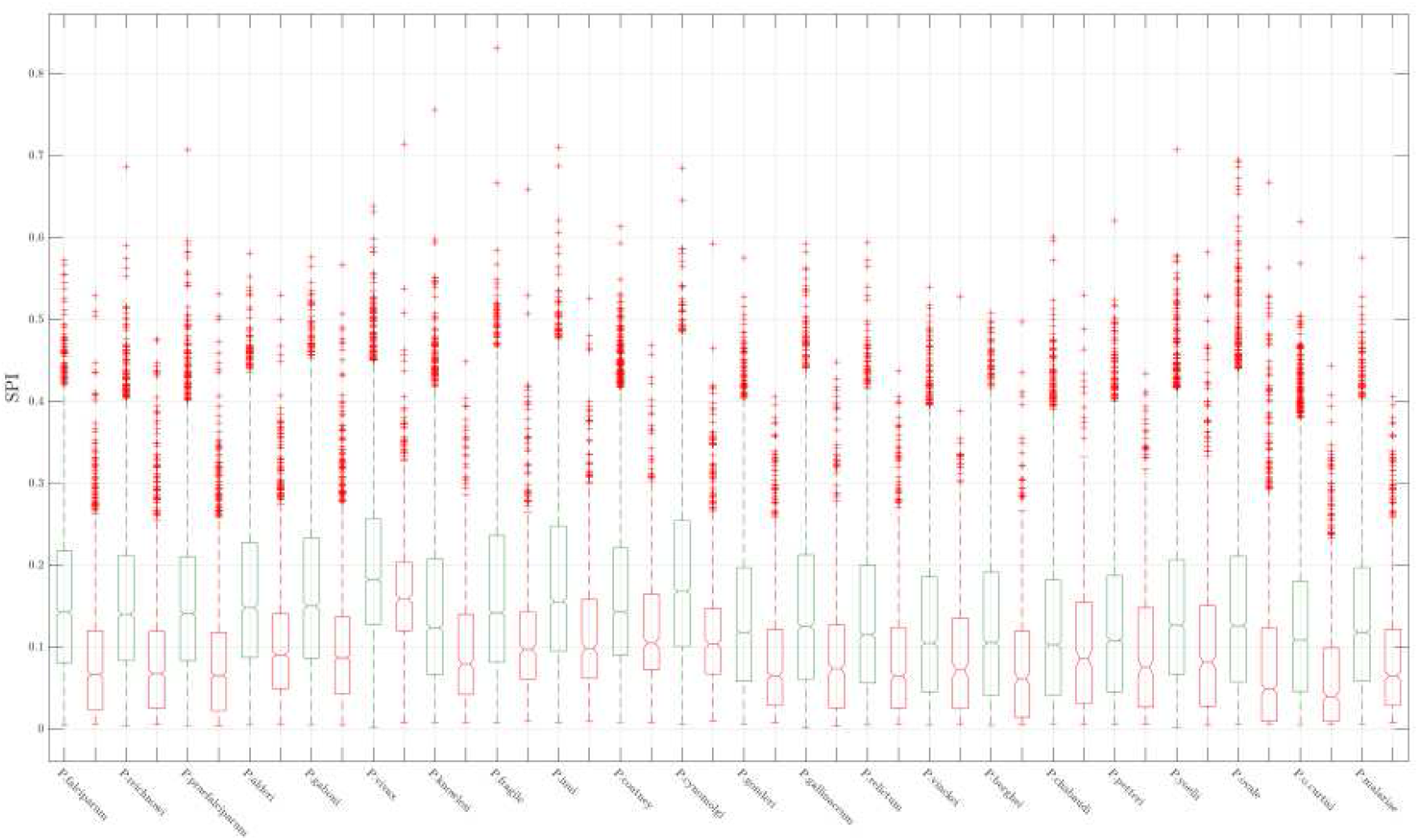
Pairwise **(Welch t-test)** comparison of the SPI distributions of LCPs and nLCPs in each parasitet LCPs are represented in red. In green nLCPs. The medians of each boxplot pair differ with 95% statistical significance. (<u>Mathworks</u>).

**Fig 9.**
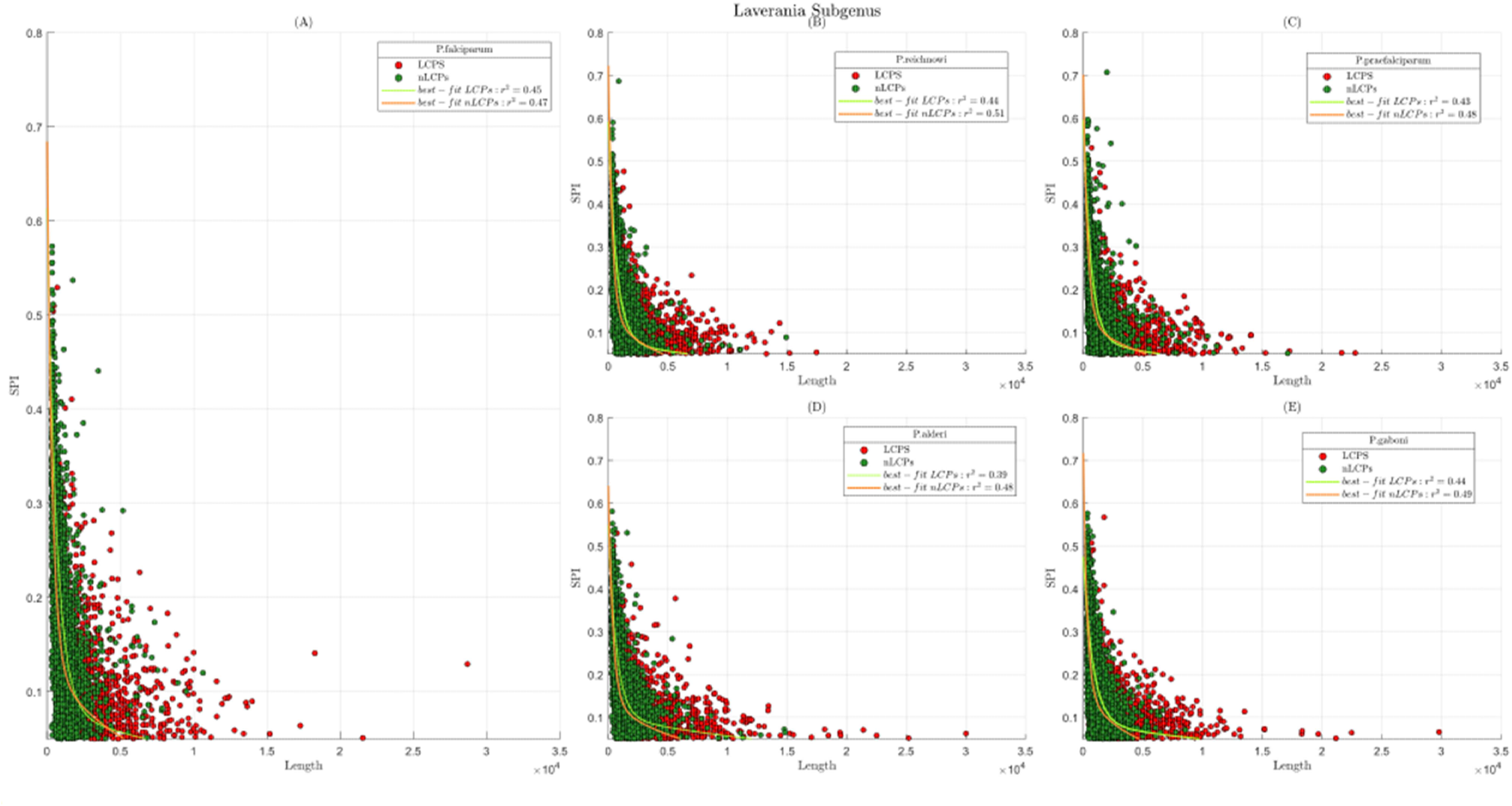
Illustration of the SPI vs length analysis performed with Laverania parasites. In red LCPs. In green nLCPs. We utilized an **a*exp(b*x) + c*exp(d*x)** model. (**A**) **P.falciparum: nLCPs’ best fit parameters (BFPs)**: **a** = 0.55 (0.4898, 0.618), **b** = −0.0028 (−0.003272, −0.002413), **c** = 0.13 (0.1085, 0.1525), **d** = −0.000183 (−0.000253, −0.0001131); **LCPs’ BFPs: a** = 0.53 (0.4595, 0.601), **b** = −0.0016 (−0.001798, −0.001371),**c** = 0.083 (0.06994, 0.09596), **d** = −7.5e-05 (−0.0001052, −4.61e-05), (**B**) **P.reichnowi**: **nLCPs’ BFPs**: **a** = 0.58 (0.5122, 0.6502), **b** = −0.0030 (−0.003502, −0.002601), **c** = 0.14 (0.1202, 0.1643),**d** = −0.000229 (−0.0002978, −0.0001602); **LCPs’ BFPs: a** = 0.51 (0.4451, 0.5775), **b** = −0.0015 (−0.001708, −0.001303), **c** = 0.08 (0.06701, 0.09343), **d** = −7.12e-05 (−0.0001015, −4.094e-05) (**C**) **P.praefalciparum**: **nLCPs’ BFPs**: **a** = 0.57 (0.5096, 0.6328), **b** = −0.0030 (−0.003265, −0.002493), **c** = 0.13 (0.1118, 0.149), **d** = −0.00018 (−0.0002396, −0.000124); **LCPs’ BFPs**: **a** = 0.51 (0.4397, 0.5836), **b** = −0.0016 (−0.00181, −0.001351), **c** = 0.084 (0.07013, 0.09811), **d** = −8.149e-05 (−0.0001133, −4.968e-05) (**D**) **P.alderi**: **nLCPs’ BFPs**: **a** = 0.14 (0.1177, 0.1604), **b** = −0.00017 (−0.0002284, −0.0001093), **c** = 0.50 (0.4461, 0.5571), **d** = −0.0027 (−0.003079, −0.00226); **LCPs’ BFPs**: **a** = 0.4437 (0.3759, 0.5114), b = −0.001489 (−0.001727, −0.001252), **c** = 0.10 (0.09086, 0.1175), **d** = −6.124e-05 (−8.405e-05, −3.842e-05) (**E**) **P.gaboni: nLCPs’ BFPs**: **a** = 0.55 (0.4769, 0.6341), **b** = −0.003121 (−0.003681, −0.00256), **c** = 0.16 (0.1342, 0.1881, **d** = −0.0002504 (−0.0003267, −0.0001741); **LCPs’ BFPs: a** = 0.39 (0.3423, 0.4314), **b** = −0.00124 (−0.001438, −0.001038), **c** = 0.090 (0.07191, 0.1039), **d** = −5.822e-05 (−8.891e-05, −2.752e-05). All the coefficients are provided with their 95% confidence bounds

### Pr2

To strengthen our confidence regarding Darwinian Pressures shaping CUB of Plasmodium species, a Parity Rule 2 (**Pr2**) analysis was performed (**Fig.10**). Once more, we calculated Pr2 scores, separately, for LCPs and nLCPs represented in red and green, respectively. Pr2 plots are provided with centroids of nLCP and LCP distributions. Laverania, Vinckeia and Haemamoeba Subgenera show distributions located near the centre of Pr2 plots even if their tails tend to diverge towards the external edges of Pr2 planes, confirming both the actions of selective pressure and mutational bias. Diversely from Vinckeia Subgenus, whose parasites have both their centroids placed in the second quadrant of Pr2 plot, Laverania and Haemamoeba Subgenera show to have LCP centroids on the first quadrant showing off, to a certain degree, a preference for A and G in the wobble codon position of their 4-fold codon families. Differently from Haemamoeba Subgenus, nLCP centroids of Laverania Parasites appear in the second quadrant of Pr2 plane with consequently a preference for C and G, in contrast to a preference for A and G reported for birds Plasmodia. Similar considerations to Haemamoeba Plasmodia can be drawn for HIPs. Simian Plasmodia show more rounded distributions, representative of their even GC content. Interestingly, remembering the nLCPs of P.berghei that distributed throughout the ENC plane and the AT rich islands of P.vivax and P.cynomolgi, these three clusters disappear in the respective Pr2 plot, underlying that, despite the differences evidenced through the ENC analysis, 4-fold codon families of these protein coding genes undergo to similar selective trends to the other nLCPs, vanishing in green clouds. Pr2 plot is characterized by a vectorial nature that allows to discriminate among various pressures driving protein coding genes in one quadrant of Pr2 plane rather than in another. Thus, the diversified selective pressures pushing on protein coding genes can, mathematically, neglect each other, making the centroids under-representative. Therefore, we operated as Forcelloni & Giansanti (2020), calculating the distributions of gene deviations from the Pr2 centre as:

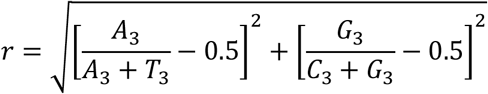

r better indicates the extent of Pr2 violations. In particular, if r = 0 the genes witness a perfect balance between natural selection and mutational bias. If r > 0 natural selection and mutational bias provide different contributions concerning a greater Pr2 violation (Forcelloni & Giansanti,2020). We compared the r distributions of nLCPs and LCPs, parasite-wise (*welch t test*) (**Fig. 11**). Likewise to the other comparisons (ENC and SPI), LCPs tend to stay closer to the parity centre with respect to the nLCPs do (p<<0.01 on average, P.fragile p<0.01) (Welch t-test). We were further intrigued by Pr2 violation and the length of protein coding genes. To unveil this information a 3D space, where (X,Y) plane is the Pr2 plane and the quote is gene length, is used. Given the difficulties of layout and the redundancies of the graphs, we show here only Laverania Plasmodia (**Fig.12**). The rest of the parasites is anyway provided in SM. We observe that longer proteins tend to cluster near the centroids that in turn do not deviate consistently from the parity centre, characteristic that is in accordance with what predicted through SPI analysis. So, LCRs are more abundant where the Pr2 deviation is smaller. Overall, Pr2 analysis sustains what we stressed through SPI analysis highlighting through 4-fold codons the relationship between mutational bias and LCRs.

**Fig 10.**
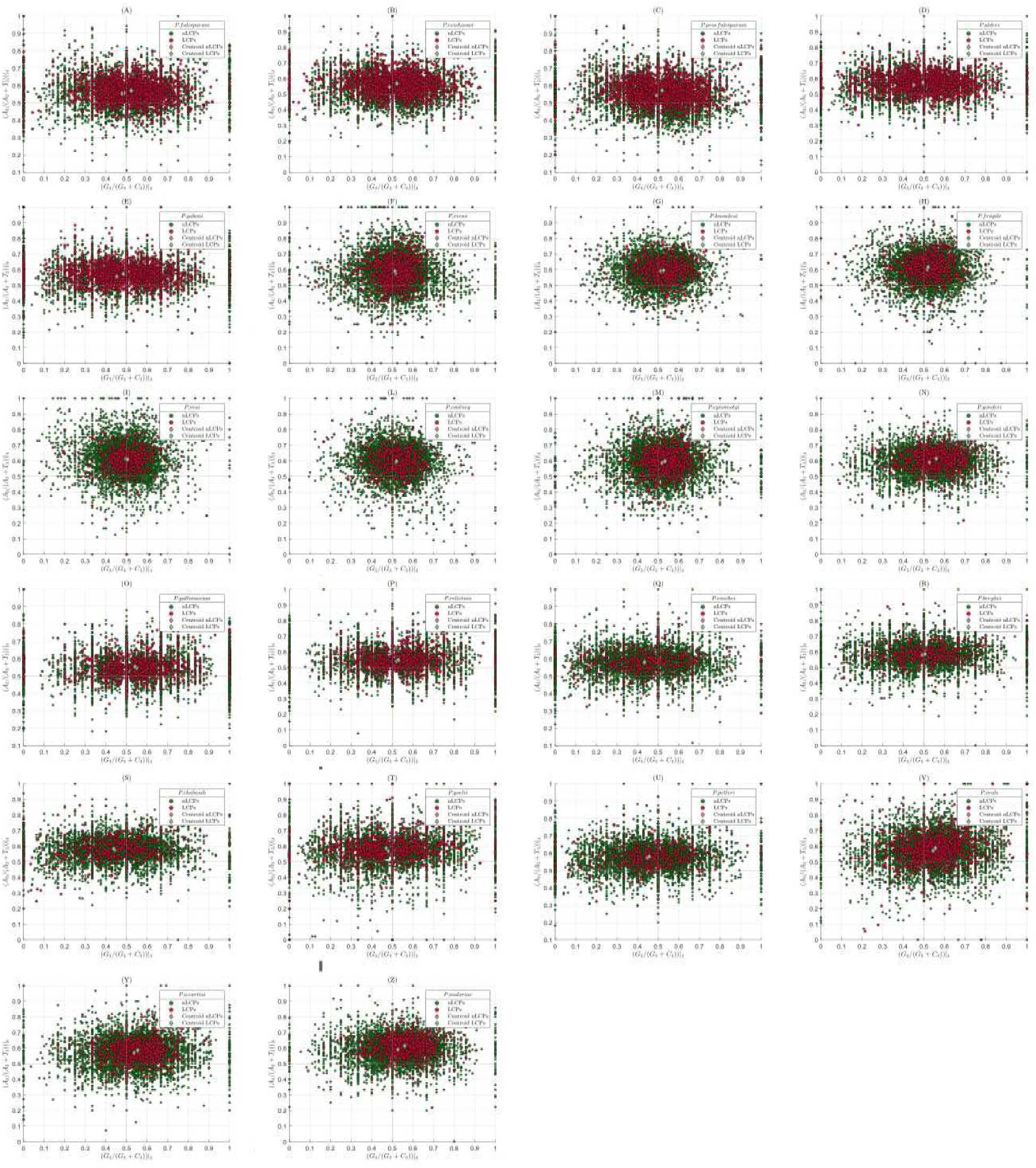
Pr2 plots. (**A, B, C, D, E**) Laverania Plasmodia; (**F, G, H, I, L, M, N**) Simian Plasmodia; (**O,P**) Haemamoeba Plasmodia;(**Q,R,S,T,U**) Vinckeia Plasmodia; (**V, Y,Z**) Human Infecting Plasmodia. Moving away from the Pr2 center returns the extent with which Parity Rule 2 is violated in a Protein Coding Sequence. The more a CDS moves away from the center, the more the contribution to the CUB can be attributed to Selective Pressure.

**Fig 11.**
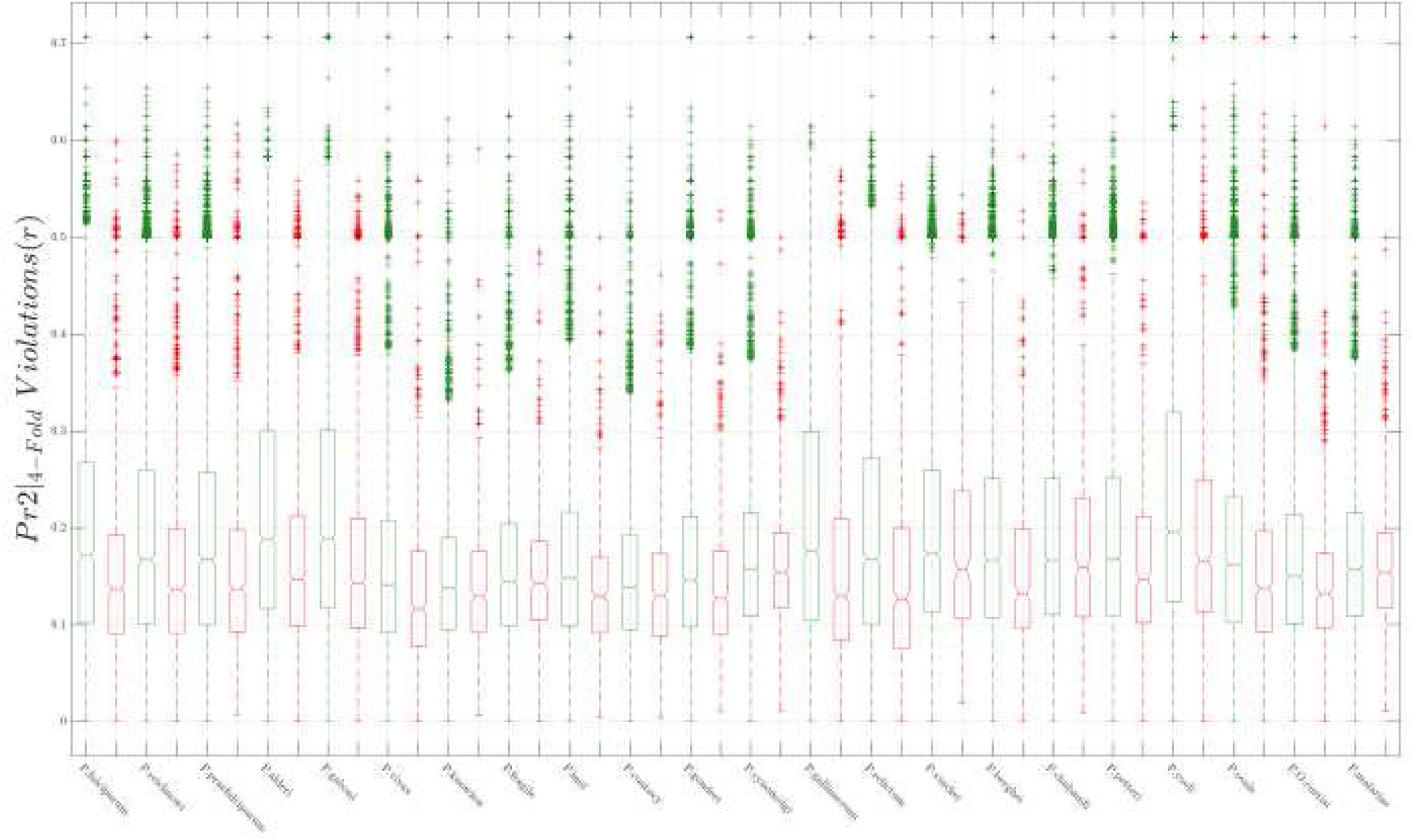
Pairwise comparison between Pr2 Violation distributions of LCPs and nLCPs represented in red and green, respectively. Medians in each pair differ with 95% confidence (<u>Mathworks</u>).

**Fig 12.**
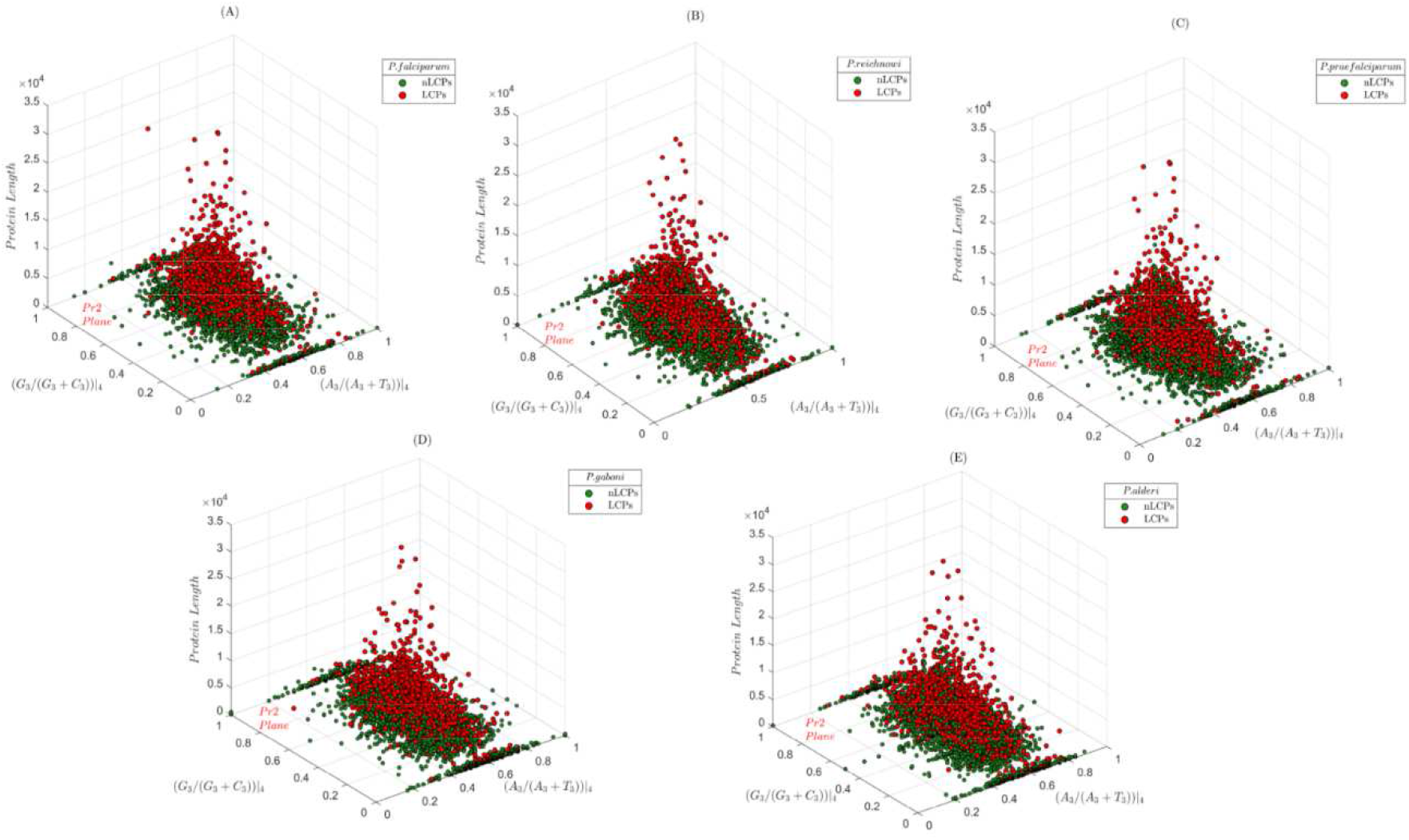
Correlation between Pr2 data and protein length. Pr2 plot is represented in the (X,Y) plane while protein length (nucleotides) is placed over the quote

## Discussion

Malaria is one of the most severe public health problems worldwide being the leading cause of death and disease in many developing countries, with 405.000 estimated deceases in 2018 (CDC). Comparative genomics is a powerful tool to unravel evolutionary changes among organisms, helping to identify the biological characteristics that give each life form its unique attributes (Nature Education). In this study, we have derived a set of information regarding the selective pressures acting on 22 Plasmodium species in order to then be able to advance a hypothesis regarding the nature of LCRs in P. falciparum.

### GC content

In line with the literature (Zilversmit et al.2010; DePristo et al.2006; Frugier et al. 2009; Filisetti et al. 2013; Pizzi & Frontali,2001; Xue & Forsdyke, 2003), the P. falciparum proteome presented low levels of Guanine and Cytosine. In this regard, the analysis has begun to emphasize a marginal relationship between genomic bias and quantity Low Complexity Regions abundance. Our statistical comparisons do not support the hypothesis that the preponderance in LCRs of the Laverania family is driven by genomic bias. In P. falciparum, Hamilton and colleagues (2016) have shown *in vitro* a pronounced excess G: C to A: T transitions finally attributing the genomic skew of these parasites to mutational bias. Assuming that similar transitions also occur in the other AT rich parasites, the simple genomic bias does not explain the extreme diversity of these organisms both in quantity of LCRs and in quantity of LCPs of which the Laverania family seem to be particularly abundant (see SM). In this sense, we agree with Chaudhry et al. (2017) in decreeing the role of genomic biases as marginal. On the other hand, we do not deny GC content has a strong influence in the choice of synonymous codons.

### RSCU and LCRs

We applied RSCU (Sharp & Li, 1987) utilizing LCRs DNA. We identify similar preference in the most used codons of the AT-rich species while Simian Plasmodia display a shuffled repertoire of RSCU values that is in line with their even GC content. Clustering analysis (Ward, 1963) produces relevant observations that are consistent with literature. It divides species into two main evolutionary lines, emphasizing a mutual difference for their G / C or A / T wobble positions. An evolutionary blueprint of the Protozoa is indeed shined back, conserving the sisterly relationship between P.cynomolgi and P.vivax (Tachibana et al. 2012; Hayakawa et al. 2008). P.gonderi is placed in the same lineage of Simian Plasmodia stressing, once more, an African origin for these parasites (Arisue et al. 2019) further emerging to undergo similar pressures in the choice of synonymous codons for their LCRs. Remarkably, the branching of the Laverania family reports consistent data from various other works (Otto et al. 2018; Silva et al. 2011; Rich et al. 2009) highlighting a common ancestry with Vinckeia (Ramiro et al. 2012) and Haemamoeba (Waters et al. 1991; Escalante et al. 1998) given the striking similarity of the CUB to their LCRs.

On the other hand, the heatmap returned a comparison between the RSCU values among all species blurring the choice within the single ones. The analysis of the RSCU normalized values illustrates for the species belonging to the Laverania, Vinckeia and Haemamoeba Subgenera, constituting what was seen through the heat map, a clear pressure for wobble positions in A or T. Similar conclusions are drawn for P.gonderi and for HIPs despite having been placed in the second evolutionary group by the clustering algorithm. The situation of the species belonging to the rest of the Simian family is different. In fact, the action of the *drift* (Bulmer,1991) is more pronounced where most of the time we do not find a stark preference between wobble positions in A / G or T / C. So LCRs of Simian Plasmodia, if compared to the others, appears under a lower selective pressure being these pressures apparently disputed in a tug of war between multiple synonymous codons.

### Complexity

We used Shannon’s Entropy, comparing the complexity of the TRRs belonging to these micro-organisms. We have shown that the LCRs of Laverania Plasmodia are composed of fewer types of codons than other parasites; we observed that, with the same length, the Laverania Plasmodia have less complex LCRs than the other species analysed here. Furthermore, the correlation between average complexity and GC content suggests that genomic bias cannot be the only contributor to the codon bias of LCRs. Among the various factors contributing to the instability (the proclivity to expand or contract) of a TRR there is the purity (Ngai & Saitou, 2016; Saitou, 2018; Gemayel et al. 2012; Legendre et al. 2007) of the nucleotide tract and its length (Gemayel et al. 2012; Legendre et al. 2007); the nature of the codon bias upstream of a repetitive sequence of amino acids (Verstrepen et al. 2005); the nucleotide composition (Gragg et al. 2002).As stated, on average, LCRs of Laverania Plasmodia appear to be shaped by a lower number of codons species as opposed to the other species which on average exhibit higher complexity: it suggests a major instability. Moreover (see Supplementary Materials), the predominance in the Laverania Subgenus belongs to the AAT codon that covers an interquartile range between 35% and 80% of the length of the LCRs. In the other species rich in Adenine and Thymine, this dominance is contested by GAA (E), GAT(D) and AAA(K) codons. Asparagine has proven to be dispensable in P.falciparum proteasome lid subunit 6 (Rpn6) (Muralidharan et al. 2010). However, re-proposing the question already posed, in a more general context by Gemayel and colleagues (2012), whether a repetitive unit is chosen for its mutability, or whether its mutability is rather rewarded for obtaining some effect on fitness, as far as asparagine is concerned, we would expect a similar pervasiveness also in the other AT rich parasites if it were for the first reason. Furthermore, observing the values of the SPI and the Pr2 violations of the LCPs of the AT-rich parasites, the LCPs of these parasites are under similar evolutionary pressure. Therefore, attributing the massive presence of asparagine in the Laverania Plasmodia to a lack of selective constraints against its diffusion does not appear correct since, if this were the case, we would expect, again, a much more abundant presence of N also in the Haemamoeba and Vinckeia Plasmodia. Hence, the LCRs appear to be selected from nucleotide level. As previously hypothesised by Dalby (2009) we infer asparagine to be under positive selection.

### Codon Bias

RSCU has a broad spectrum of maximum values which, however, may not reflect a predominant use of a family of codons which, despite high RSCU values, may be little used. For this reason, we wanted to quantitatively study the codon composition of LCRs. The observation of the bar charts allows to consider some important aspects of the composition of the LCRs. The first question concerns the species of codons most used, where the preponderance of asparagine (mostly encoded by AAT) is particularly accentuated in the Laverania family. In fact, the codons most present in general in all species are E, S, D and R which represent the most common substitutions for N and K (<u>NCBI-N</u>, <u>NCBI - K</u>). NCBI tables stress the similarity of the chemical-physical characteristics of these amino acids. Although a more in-depth study is necessary to better visualize the composition of LC runs, we observe a certain tendency in all species to preserve the CUB of these *loci* since amino acidic changes appear to be primarily conservative, *i.e.* that the properties of the polypeptide do not drastically change (Strachan et al. 2020). Noteworthy, LCPs of the Simian Plasmodia lie in the central part of the ENC plots which denotes a higher GC3 content. In spite of this, it is not uncommon to observe that the linear extension of the LCRs of Simian spp. is composed of codons rich in Adenine as is the case of GAA (E) which rivals its synonym GAG (see Supplementary Materials) and in general as for other codons going against the GC pressure which appears to press on their proteomes.

### Protein Length

We observed that LCPs are longer *de facto* than nLCPs. This implies that the LCRs are preferentially inserted into the longest proteins of each parasite. Pizzi E. and Frontali C. reported how some of the P. falciparum proteins emerged to be longer than the orthologues found in other organisms (Pizzi & Frontali, 2001) by virtue of the presence of LCRs as retraced by Xue and Forsdyke (2003) that report how these are placed between domains to which a specific function can be ascribed. Therefore, we hypothesize that longer proteins are selected to host LCRs because there is less chance of disrupting a functional domain, by squeezing them into interdomains. This would also explain the positive correlations between protein length and LCRs abundance found in each parasite (**Tab.1**). It is not clear whether recombination or replication slippage is the predominant mechanism. The lack of consensus in distinguishing between micro and minisatellite regions (Guy-Frank et al. 2008) makes it difficult to propose a generalized hypothesis discriminating between the two mechanisms, where in general, replication slippage is suggested for minisatellites (Guy-Frank & Paques, 2000) and recombination like phenomena are suggested for minisatellite regions (Kokoska et al. 1998) and less complex LCRs (Ellegren,2004). Notably, in P. falciparum, Zilvermit et al. (2010) suggest unequal crossing over events for GC rich LCRs that can be luckily be extended to the other Laverania spp. Overall, more evidence is needed to decree with greater reliability that LCRs space functional domains without their disruption, that is left for future research.

### Darwinian Pressures

#### Codon Bias and Gene Expression

To cope with Darwinian selective pressures, we relied on the Effective Number of Codons (Wright, 1996) using the improved version of Sun et al (2012), SPI, and PR2 (Sueoka, 1995; Sueoka & Kawanishi, 1999). These analyses consistently apply to each proteome stressing different equilibria between selective pressure and mutational bias regarding both LCPs and nLCPs. Overall, the set of these tools indicates that the CUB of LCPs is more strongly influenced by mutational bias rather than by selective pressure and that the contribution of the latter tends to decrease with the length of the protein. Optimal codon bias is often attributed to high levels of gene expression (Bulmer, 1991; Sharp & Li, 1986). Furthermore, it is assumed that a shorter protein requires less time to fold (Williamson, 2017) and that it must require a smaller number of molecular chaperones (Lipman et al 2002) for their surveillance function to be performed (Daniyan et al. 2019) with less space reserved for their folding ultimately avoiding aberrant interactions with other cellular components (Williamson, 2017; Hartl & Hartl, 2002). This would suggest better translation efficiency of nLCPs and higher expression. Duret & Mouchiroud (1999) showed this in *C.elegans* where as selective pressure (protein length) decreases (increases) gene expression increases (decreases) as well. However, transcriptomic examinations have shown that gene expression in P. falciparum is substantially different between *field isolates* and *in vitro* cultured samples, where this variability is associated with Copy Number Variations (Mackinnon et al. 2009). Other studies indicated that gene expression in P.falciparum varies according to the applied pharmacological stressor (Hu et al. 2009). Moreover, *in vitro* proteomics data show how the amount of proteins expressed in P. falciparum follows a very precise time course (Foth et al. 2011). A similar trend was also observed in P.vivax (Bozdech et al. 2008). This suggests that the gene expression of these parasites may vary depending on external factors and depending on the stage of the life cycle in which they are found. Therefore, the association of the two gene categories (nLCPs and LCPs) to a lower or higher gene expression is controversial and not so easily attributable.

#### SPI and Pr2 violations: Positive and Negative Selective Pressure

The SPI is intrinsically linked to the nature of the Effective Number of Codons (Wright, 1996) and it is suggested to be used coupled with the improved version of Sun and colleagues (2012) or any other improved version of the ENC such as that of Fuglsang (2006). MT1 revealed the SPI to be in line with the purpose for which it was conceived. Compared to Laverania Plasmodia, Simian spp. show a more heterogeneous codon bias (their ENC values are larger, see **Fig. 7**). LPCs of Simian Plasmodia show off the highest SPI values, underlining that a higher number of codons used does not necessarily identify a lower selective pressure. We agree with Gajbhiye and colleagues (2017) in decreeing a greater selective pressure on the proteins of P.vivax compared to those of P. falciparum, to which the other members of its family are also added. Anyway, SPI it is not able to distinguish between positive and negative pressures but is only able to establish what the divergence from Wright’s Theoretical Curve is, studying how closely the CUB of a gene approximates an extreme Bias situation (20 codons for 20 amino acids). Similarly, Pr2 (Sueoka, 1995; Sueoka & Kawanishi, 1999) allows to understand what the shift from Parity Rule 2 is, returning, like SPI, a departure from a situation of equilibrium. The genomes of malaria parasites suffer, in respect of many genes, an apparent lack of orthologues (Gardner et al. 2002; Hu et al. 2009) which causes a complication for the application of methods such as, *e.g*, Nei and Gojobori’s (1986). Addressing these problems is therefore left to future efforts.

#### Selective Pressure on LCPs and LCRs abundance

The negative correlation between selective pressure and protein length, particularly in the Laverania Plasmodia, hides information. In fact, looking to the correlation between protein length and LCR abundance (**Tab. 1**), it can be observed that proteins with a minor deviation from Wright’s Theoretical Curve and with a minor violation of Pr2 contain more Low Complexity Regions. As the complexity of LCRs increases, a functional role is suggested (Zilversmit et al. 2010). However, as instance, contrary to the tendency of long E runs to generate distortions in a polypeptide chain (Karlin et al. 2001), *in vitro* assays highlight how the areas rich in glutamate (E) in P.falciparum help drive the immune response away from the functional domains of the proteins thus evading the host’s immune system (Hou et al. 2020).Therefore, hypothesising a functional role even for pure or almost pure amino acid run it is possible and classifying pure LCRs as a neutral insertion could be not correct in all the circumstances. A hypothesis of interest is that of functional amyloid, which states that many organisms have evolved to take advantage of the potential tendency of some residues to form this type of fibrils (Fowler et al. 2007). A shared opinion is that intracellular parasites possess simplified genomes to adapt to the host (Daniyan et al. 2019). These simplifications are believed to lead to the expression of mutated proteins prone to aggregation (Daniyan et al. 2019). Despite the tendency of asparagine to form amyloid fibrils (Halfmann et al. 2011) it is reported that there is no *in vivo* evidence of such aggregates in P. falciparum (Muralidharan et al. 2010) which suggests that the parasite has learned to take advantage or at least to manage this amino acid. Fairly recent studies (Halfmann et al. 2011) have clearly highlighted how LCRs rich in asparagine, in *yeast*, reduce the toxicity of proteins. Given the negative correlation between selective pressure and LCRs abundance, it would be interesting to look for a possible concomitance of detrimental polymorphisms and the presence of asparagine, which would suggest a buffer role for these amino acid regions. Overall, these correlations are puzzling and open an exceptionally large number of avenues for new hypothesis and research.

#### Criticisms, Possible Refinements and Future Addresses

An important concept that is necessary to point, concerns SEG parameters. In fact, standard parameters allow to identify regions strongly polarized towards a certain species of amino acids whilst allowing to find LCRs with a more heterogeneous repertoire of amino acids (Trilla & Albà, 2012). On the other hand, enlarging the window through which SEG is set introduces a reduction in the LCRs that are identified, where smaller windows (such as W=6) allow to observe a larger number of LCRs (Batistuzzi et al. 2016). Comparing the statistics provided by our work with some others (Chaudhry et al. 2018), on a qualitative level, the identified amino acids are not starkly different from those of Chaudhry et al. (2018) and the proportions between nLCPs and LCPs are consistent with those found by them (see SM). However, the critical sense suggests validating the subjectivity with which SEG is triggered, finding a trade-off between the classical approach with standard parameters and a more refined one such as that proposed by Batistuzzi and colleagues (2016) through by which to identify any artificial classification in LCPs and nLCPs stratification. Likewise, it is important to point that GC content often aggravates genome assemblies (Chen et al. 2013). Even though what is reported in the Supplementary Materials confers a certain robustness to our observations, finding similar results for SPI and protein length in other genome assemblies (same species different data), the risk is not absent. Shannon’s Entropy, as used in this work, made us understand important characteristics regarding the composition of LCRs. However, making this implementation more general and more widely usable is left to future work. Finally, a functional enrichment in GO terms could indicate the phenotype associated with the proteins where the LCRs are inserted, providing with more clues the rationale to infer the directionality Darwinian Selective Pressure.

## Conclusion

Overall, in this study we tackled the proteome analysis of 22 plasmodia providing important insights into the pervasiveness of Asparagine Low Complexity Regions. Extending the present research to other *Apicomplexa* would lead to a deepened knowledge of the biology of these parasites, reaching a better understanding of their evolutionary success.

## Contribution

C.A. and G.A. conceived the study. C.A. conducted the analyses. C.A. and G.A. wrote the main manuscript. C.A. A.G. and F.S. read and approved the final manuscript.

