## Supplementary Materials for "Evolutionary Pressures and Codon Bias in Low Complexity Regions of Plasmodia Parasites"

**Introduction**

We have collected some of the results of the main body of work and other analyses. This, due to the excessive number of graphics and information. In the first paragraph we give the number of protein sets analysed in our work. Next, the relationships between protein length vs SPI and protein length vs Pr2. We have also studied, although a more in-depth analysis is postponed to future works, also other assemblies relating to the species considered in the main body of the article, which corroborates what was previously observed.

**Tab. SM1**

| **Organism** | **nLCPs** | **LCPs** | **% LCPs** | **% nLCPs** |
| --- | --- | --- | --- | --- |
| P.falciparum | **3638** | **1877** | **34** | **66** |
| P.reichnowi | **3892** | **1876** | **32** | **68** |
| P.praefalciparum | **4185** | **1844** | **30** | **70** |
| P.alderi | **3609** | **1805** | **34** | **66** |
| P.gaboni | **3661** | **1784** | **33** | **67** |
| P.vivax | **5616** | **848** | **13** | **87** |
| P.knowlesi | **4523** | **586** | **12** | **88** |
| P.fragile | **5008** | **664** | **14** | **86** |
| P.inui | **5315** | **517** | **9** | **91** |
| P.coatney | **5008** | **508** | **10** | **90** |
| P.cynomolgi | **5012** | **704** | **13** | **87** |
| P.gonderi | **5131** | **786** | **14** | **86** |
| P.vinckei | **4558** | **396** | **8** | **92** |
| P.petteri | **4773** | **387** | **8** | **92** |
| P.berghei | **4596** | **422** | **9** | **91** |
| P.yoeli | **6984** | **877** | **11** | **89** |
| P.chabaudi | **4762** | **409** | **8** | **92** |
| P.gallinaceum | **4430** | **875** | **17** | **83** |
| P.relictum | **4458** | **791** | **16** | **84** |
| P.malariae | **5131** | **786** | **14** | **86** |
| P.ovale | **7457** | **1124** | **14** | **86** |
| P.o.curtisi | **6386** | **776** | **11** | **9** |

**SPI and Length correlations**

We report the remaining fits between SPI and protein length. As previously mentioned, proteins are placed in a plane where SPI is the ordinate and protein length is the abscissa. The parameters of the best fit are reported in the caption of each image with the respective 95% confidence bounds. We point that LCPs are depleted of their LCRs.

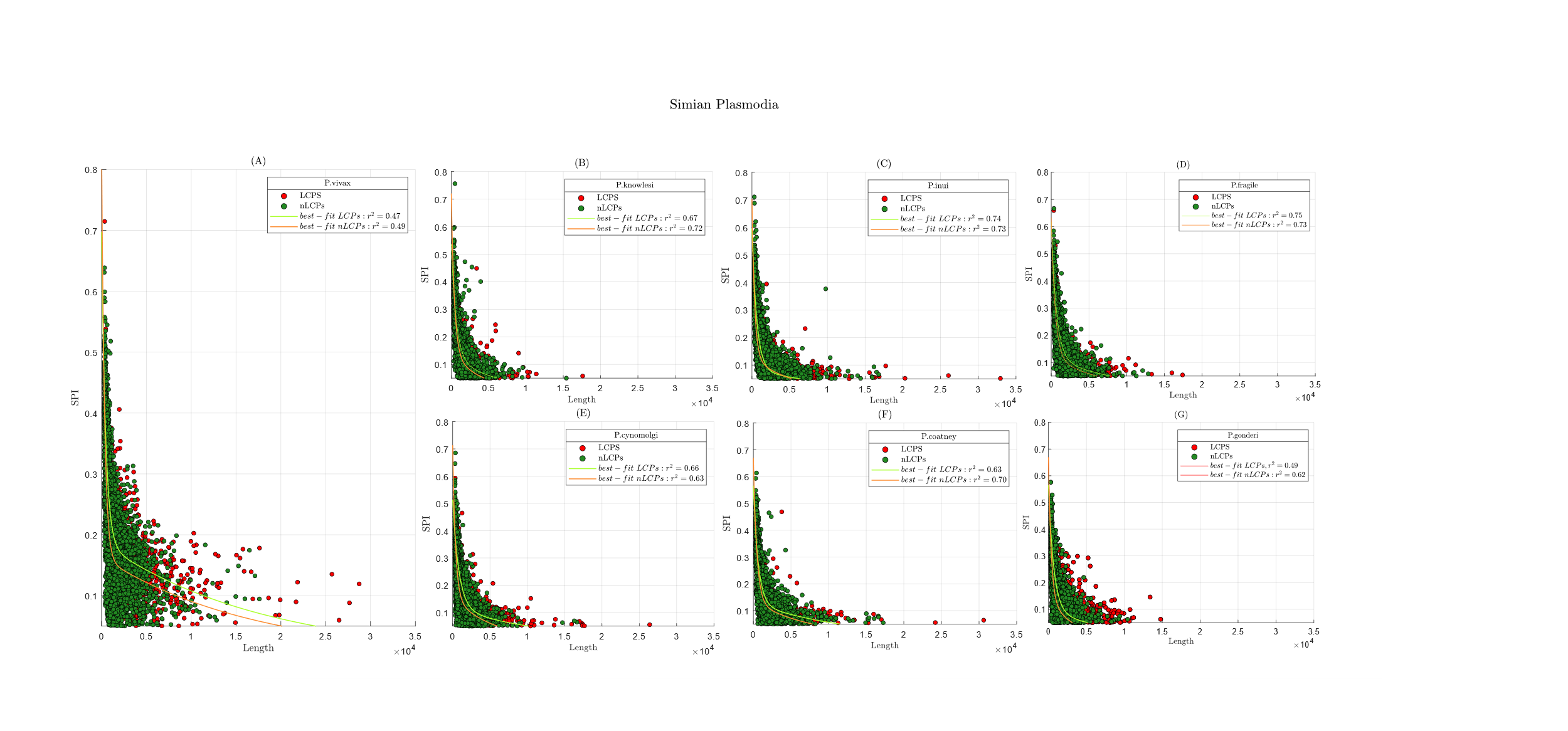

**Fig. SM1** Illustration of the SPI vs length analysis performed. **Simian Plasmodia**. In red LCPs. In green nLCPs. We utilized an **a*exp(b*x) + c*exp(d*x)** model. (**A**) **P.vivax** : **nLCPs’ BFPs**: **a** = 0.693,**b** = -0.003144 (-0.003395, -0.002892),**c** = 0.1634 (0.1554, 0.1713), **d** = -5.885e-05 (-7.501e-05, -4.27e-05); **LCPs’ BFPs :** **a** = 0.5099 (0.3856, 0.6343), **b** = -0.002237 (-0.002711, -0.001763), **c** = 0.1873 (0.1743, 0.2004),**d** = -5.528e-05 (-6.898e-05, -4.158e-05); (**B) P.knowlesi: nLCPs’ BFPs:** **a** = 0.5777 (0.5412, 0.6142),**b** =	-0.002449 (-0.002679, -0.00222),**c** =	0.143 (0.1262, 0.1598),**d** = -0.0002384 (-0.0002797, -0.0001972); **LCPs’ BFPs:** **a** = 0.4109 (0.3356, 0.4863);**b** =	-0.001546 (-0.001959, -0.001134);**c** = 0.1263 (0.09275, 0.1599); **d** = -0.0001503 (-0.0002062, -9.441e-05) (**C**) **P.inui: nLCPs’ BFPs:** **a** = 0.5204 (0.4945, 0.5463),**b** = -0.002269 (-0.002458, -0.002081), **c** = 0.1536 (0.1404, 0.1669),**d** = -0.0001667 (-0.0001944, -0.0001389); **LCPs’ BFPs:** **a** = 0.487 (0.4257, 0.5483);b = -0.001472 (-0.001732, -0.001212); **c** = 0.1217 (0.1005, 0.1429); **d** = -8.254e-05 (-0.0001139, -5.121e-05) (**D**) **P.fragile:** **nLCPs’ BFPs:** a = 0.5091 (0.4875, 0.5306),**b** = -0.001951 (-0.00211, -0.001792),**c** = 0.1244 (0.1094, 0.1394),**d** = -0.0001508 (-0.0001864, -0.0001153);**LCPs BFPs:** **a** = 0.4966 (0.4413, 0.5518);b =	-0.00148 (-0.001716, -0.001244);c =	0.1208 (0.09831, 0.1434);**d** = -0.0001117 (-0.0001498, -7.356e-05) (**E**) **P.cynomolgi: nLCPs’ BFPs:** **a** = 0.5547 (0.515, 0.5945); **b** = -0.002657 (-0.002931, -0.002383);**c** =	0.1587 (0.1421, 0.1754); **d** = -0.0001927 (-0.0002324, -0.0001531); **LCPs’ BFPs**: **a** = 0.3931 (0.3438, 0.4425); **b** = -0.001217 (-0.00148, -0.0009531); **c** = 0.1196 (0.09252, 0.1467);**d** = -8.583e-05 (-0.0001256, -4.607e-05); (**F**) **P.coatney: nLCPs’ BFPs:** **a** = 0.5345 (0.5071, 0.5618); **b** = -0.002186 (-0.00235, -0.002022); **c** = 0.1359 (0.1248, 0.1469); **d** = -0.0001276 (-0.0001519, -0.0001032); **LCPs’ BFPs:** **a** = 0.4368 (0.3572, 0.5164); **b** = -0.001576 (-0.001934, -0.001218);**c** = 0.1295 (0.1068, 0.1521);**d** = -8.29e-05 (-0.0001161, -4.967e-05);(**G**) **P.gonderi: nLCPs’ BFPs:** a = 0.5274 (0.4955, 0.5593);**b** =	-0.002325 (-0.002555, -0.002096);c = 0.1045 (0.08719, 0.1218);**d** = -0.0002015 (-0.0002581, -0.000145); **LCPs’ BFPs:** **a** = 0.09095 (0.0647, 0.1172);**b** = -8.408e-05 (-0.0001303, -3.786e-05); **c** = 0.3517 (0.29, 0.4133);**d** = -0.001178 (-0.001489, -0.0008678)

**Fig SM2** Illustration of the SPI vs length analysis: Vinckeia Subgenus. In red LCPs. In green nLCPs. We utilized an **a*exp(b*x) + c*exp(d*x)** model. (**A**)**P.vinckei**: **nLCPs’ BFPs**: **a** = 0.5126 (0.4668, 0.5584);**b** = -0.002634 (-0.002983, -0.002285);**c** = 0.1328 (0.111, 0.1547); **d** = -0.0002906 (-0.0003534, -0.0002279); **LCPs’ BFPs**: **a** = 0.3836 (0.3109, 0.4564); **b** = -0.00114 (-0.001523, -0.0007564);**c** = 0.06723 (0.02446, 0.11);**d** = -7.651e-05 (-0.0001819, 2.889e-05)(B) **P.petteri**: **nLCPs’ BFPs**: **a** = 0.5187 (0.4681, 0.5692);**b** = -0.00286 (-0.003264, -0.002455);**c** =	0.1505 (0.1266, 0.1744);**d** = -0.0003487 (-0.0004137, -0.0002837); **LCPs’ BFPs**: **a** = 0.4662 (0.3759, 0.5566);**b** =	-0.001456 (-0.001934, -0.0009777);**c** = 0.09892 (0.04869, 0.1492);**d** =	-0.000133 (-0.0002368, -2.931e-05)(**B**) **P.petteri**: **nLCPs’ BFPs**: **a** = 0.5187 (0.4681, 0.5692);**b** = -0.00286 (-0.003264, -0.002455); **c** =	0.1505 (0.1266, 0.1744);**d** = -0.0003487 (-0.0004137, -0.0002837); **LCPs’ BFPs**: **a** = 0.4662 (0.3759, 0.5566);**b** =	-0.001456 (-0.001934, -0.0009777);**c** = 0.09892 (0.04869, 0.1492);**d** =	-0.000133 (-0.0002368, -2.931e-05)(C) **P.berghei**: **nLCPs’ BFPs**: **a** = 0.4867 (0.4487, 0.5246);**b** =	-0.002367 (-0.002687, -0.002047);**c** = 0.1186 (0.09422, 0.143);**d** = -0.0002711 (-0.000344, -0.0001981); **LCPs’ BFPs**: **a** = 0.3692 (0.2965, 0.4418);**b** =	-0.001201 (-0.001632, -0.000769);**c** = 0.07777 (0.02848, 0.1271);**d** = -0.0001116 (-0.000224, 7.375e-07)(**D**) **P.chabaudi:** **nLCPs’ BFPs**: **a** = 0.5029 (0.4617, 0.544);**b** =	-0.002531 (-0.002871, -0.002191);**c** =	0.1274 (0.1033, 0.1515);**d** = -0.0003026 (-0.0003733, -0.000232); **LCPs’ BFPs**: **a** = 0.1144 (0.04105, 0.1878);**b** = -0.0001504 (-0.0002784, -2.231e-05); **c** =	0.3763 (0.2807, 0.472);**d** =	-0.001309 (-0.001933, -0.000685)(**E**) **P.yoel**i: **nLCPs’ BFPs**: **a** = 0.5966 (0.5264, 0.6668); **b** = -0.003217 (-0.003677, -0.002757);**c** = 0.146 (0.1244, 0.1676);**d** = -0.0002758 (-0.0003421, -0.0002094); **LCPs’ BFPs**: **a** = 0.103 (0.06251, 0.1436);**b** = -0.0001363 (-0.0002302, -4.244e-05);**c** = 0.4419 (0.3628, 0.5211);**d** =	-0.001677 (-0.002166, -0.001188)

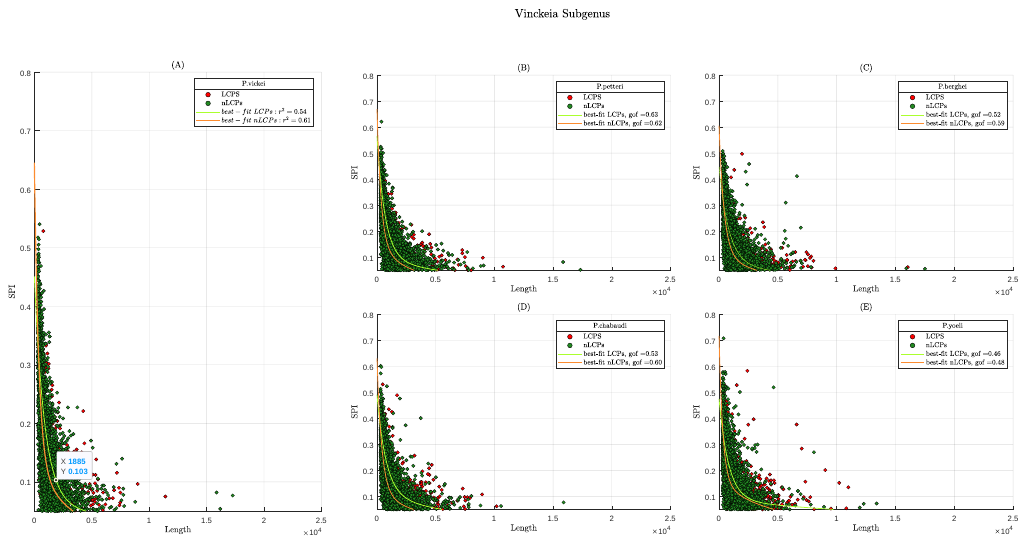

**Figure SM3** Illustration of the SPI vs length analysis: **Haemamoeba Subgenus**. In red LCPs. In green nLCPs. We utilized an **a*exp(b*x) + c*exp(d*x) model**.(**A**) **P.gallinaceum**: **nLCPs’ BFPs**: **a** = 0.4397 (0.4122, 0.4671);**b** =	-0.001813 (-0.002035, -0.001592);**c** = 0.08773 (0.06896, 0.1065);**d** = -9.849e-05 (-0.0001577, -3.929e-05**); LCPs’ BFPs**: **a** = 0.2888 (0.2395, 0.3382);**b** = -0.0009849 (-0.001318, -0.0006523);**c** = 0.09045 (0.05071, 0.1302);**d** = -9.784e-05 (-0.0001656, -3.007e-05) (**B**)**P.relictum**: **nLCPs’ BFPs**: **a** =	0.4497 (0.4149, 0.4846);**b** =	-0.002153 (-0.002446, -0.001861);c =	0.107 (0.08593, 0.128);**d** = -0.0001751 (-0.0002389, -0.0001114); **LCPs BFPs**: **a** = 0.3254 (0.2635, 0.3873);**b** =	-0.001103 (-0.001484, -0.0007212);**c** = 0.09955 (0.05562, 0.1435);**d** = -0.0001262 (-0.0002002, -5.233e-05)

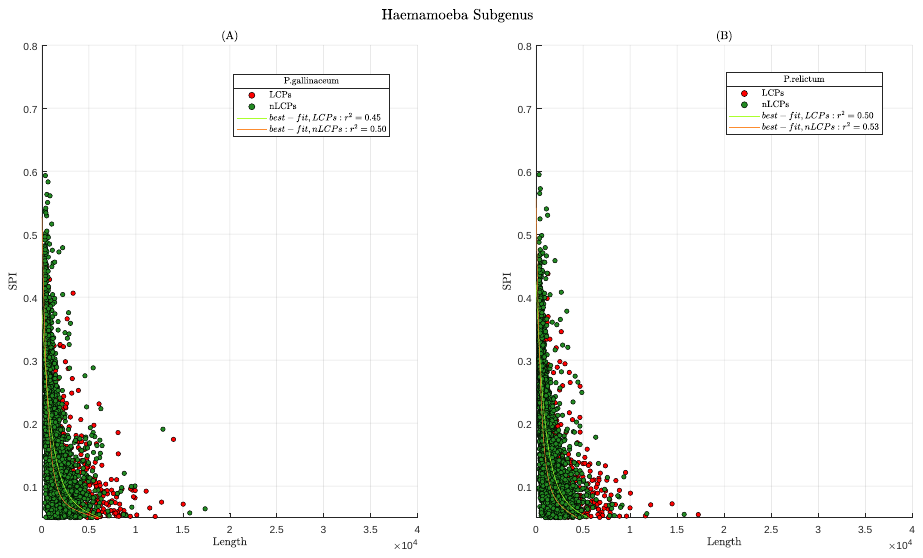

**Fig. SM4** Illustration of the SPI vs length analysis: **Human Plasmodia**. In red LCPs. In green nLCPs. We utilized an **a*exp(b*x) + c*exp(d*x) model (A) P.ovale**: **nLCPs’ BFPs**: **a** = 0.5333 (0.5072, 0.5593);**b** =	-0.002377 (-0.002674, -0.002081);**c** =	0.1113 (0.07951, 0.1431);**d** = -0.0003369 (-0.0004467, -0.0002271); **LCPs BFPs**: **a** =	0.6446 (0.5291, 0.76);**b** =	-0.002562 (-0.003172, -0.001952);**c** =	0.1512 (0.1118, 0.1907); **d** =	-0.000305 (-0.0003878, -0.0002223);(**B**)**P.o.curtisi**: **nLCPs’ BFPs**: **a** = 0.5634 (0.5306, 0.5962);**b** =	-0.002455 (-0.002723, -0.002187);**c** =	0.1033 (0.07931, 0.1274);**d** = -0.0003027 (-0.0003899, -0.0002155); **LCPs BFPs**: **a** =	0.4412 (0.3745, 0.508);**b** =	-0.001197 (-0.001451, -0.0009428);**c** =	0.05116 (0.02517, 0.07716);**d** = -8.093e-05 (-0.0001631, 1.292e-06).(**C**)**P.malariae: nLCPs’ BFPs**: **a** = 0.5274 (0.4955, 0.5593);**b** =	-0.002325 (-0.002555, -0.002096);**c** = 0.1045 (0.08719, 0.1218);**d** = -0.0002015 (-0.0002581, -0.000145); **LCPs’ BFPs:** **a** = 0.09095 (0.0647, 0.1172);**b** = -8.408e-05 (-0.0001303, -3.786e-05);**c** =	0.3517 (0.29, 0.4133);**d** =	-0.001178 (-0.001489, -0.0008678)

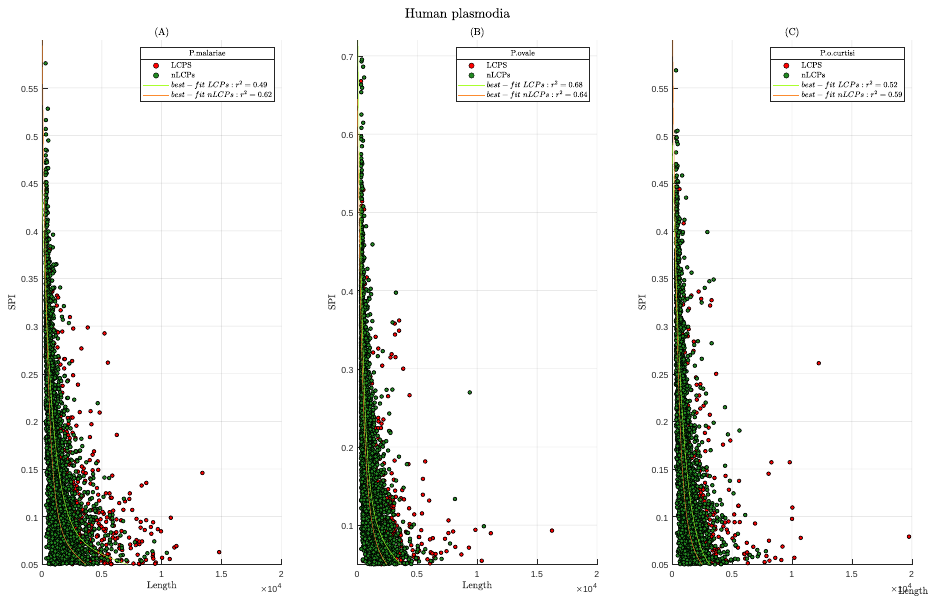

**Pr2 3D Plots**

Here we report the Pr2 correlations with protein length. Following, *Simian Plasmodia*, *Vinckeia Subgenu*s, *Haemamoeba Subgenus* and HIPs.

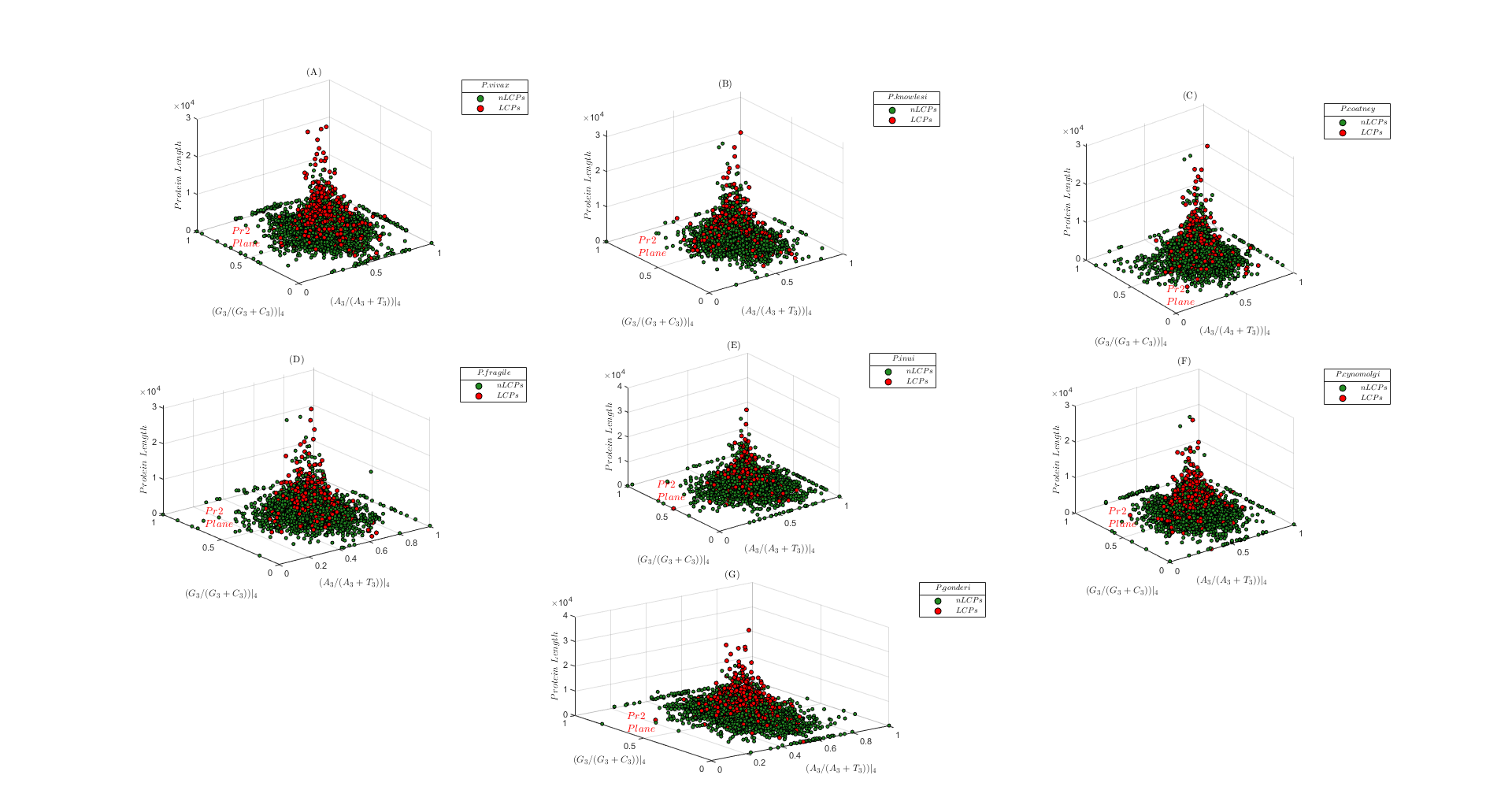

**Fig. SM5** Simian Plasmodia (**A**) P.vivax; (**B**) P.knowlesi; (**C**) P.coatney; (**D**) P.fragile; (**E**) P.inui; (**F**) P.cynomolgi; (**G**) P.gonderi

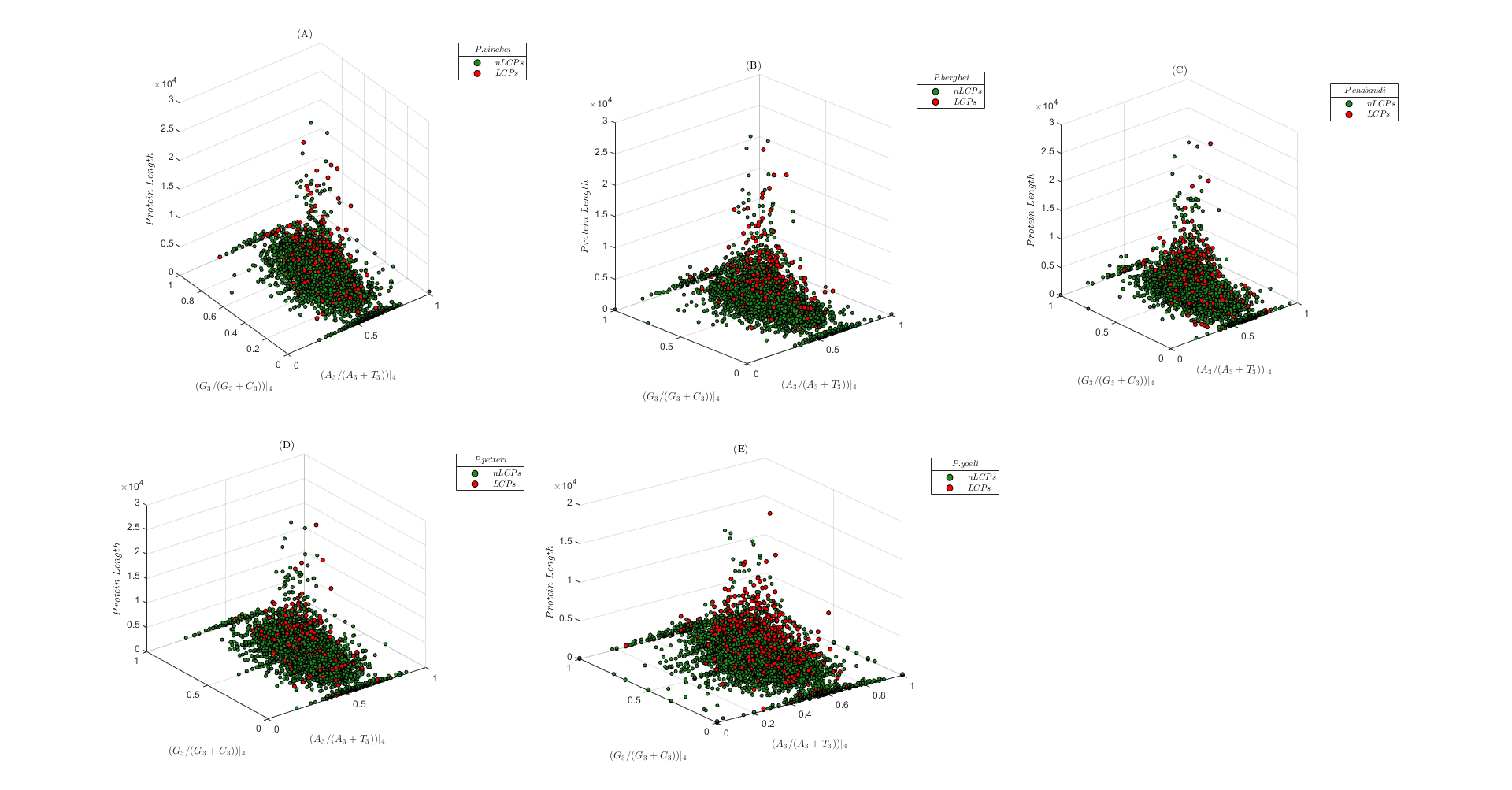

**Fig.SM6** Vinckeia Subgenus. (**A**) P.vinckei; (**B**) P.berghei ; (**C**) P.chabaudi ; (**D**) P.petteri; (**E**) P.yoelii

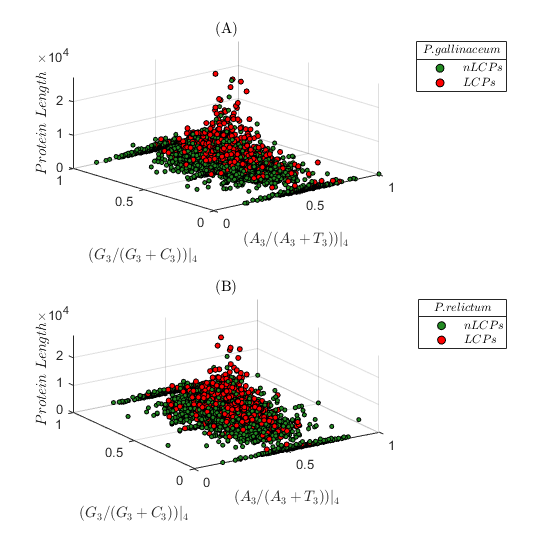

**Fig. SM7** Haemamoeba Subgenus (A) P.gallinaceum; (B) P.relictum

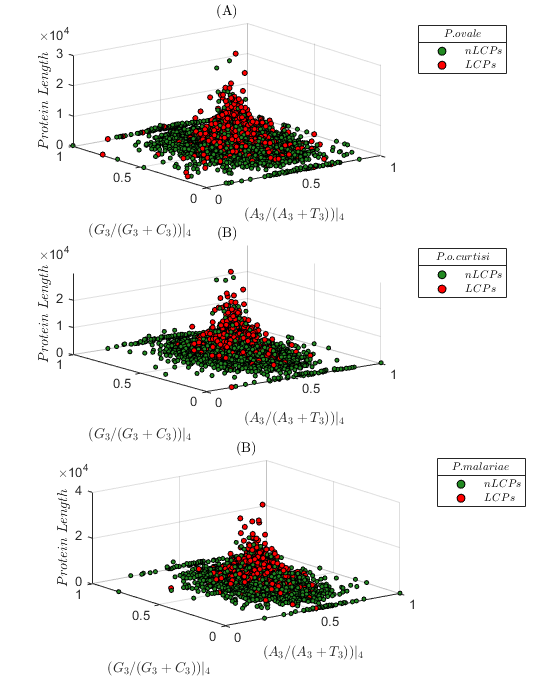

**Fig. SM7** **(A**) P.ovale (**B**) P.o.curtisi (**C**) P.malariae

**RSCU comparation**

Although other visualization strategies can certainly provide further insights, the division of 6-fold codon families eliminate the blurring from codon preference we find in the original version of RSCU.

**Fig.SM8** (**A**) Original RSCU by Sharp & Li (**1987**); (**B**) Modified Version of RSCU

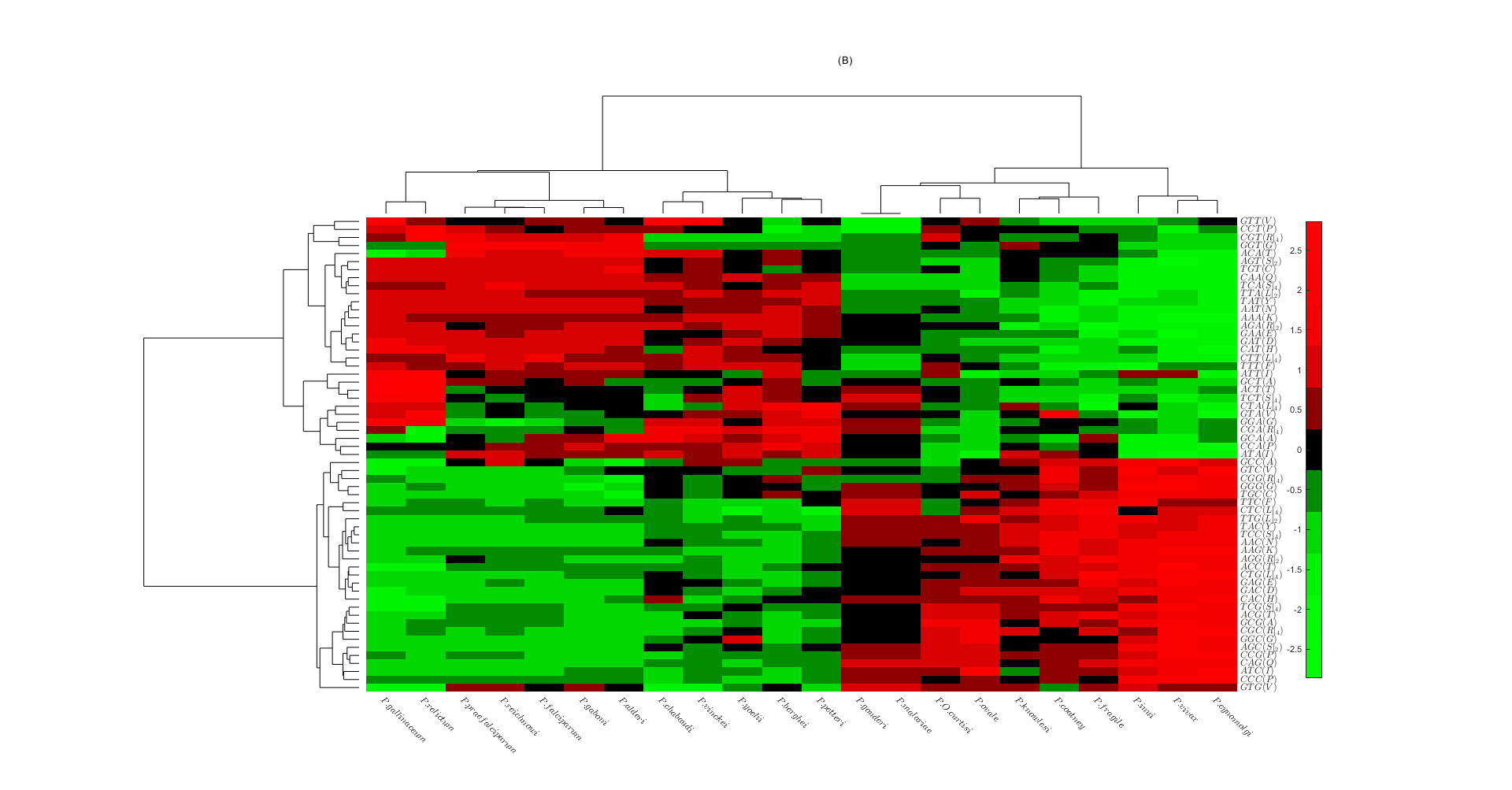

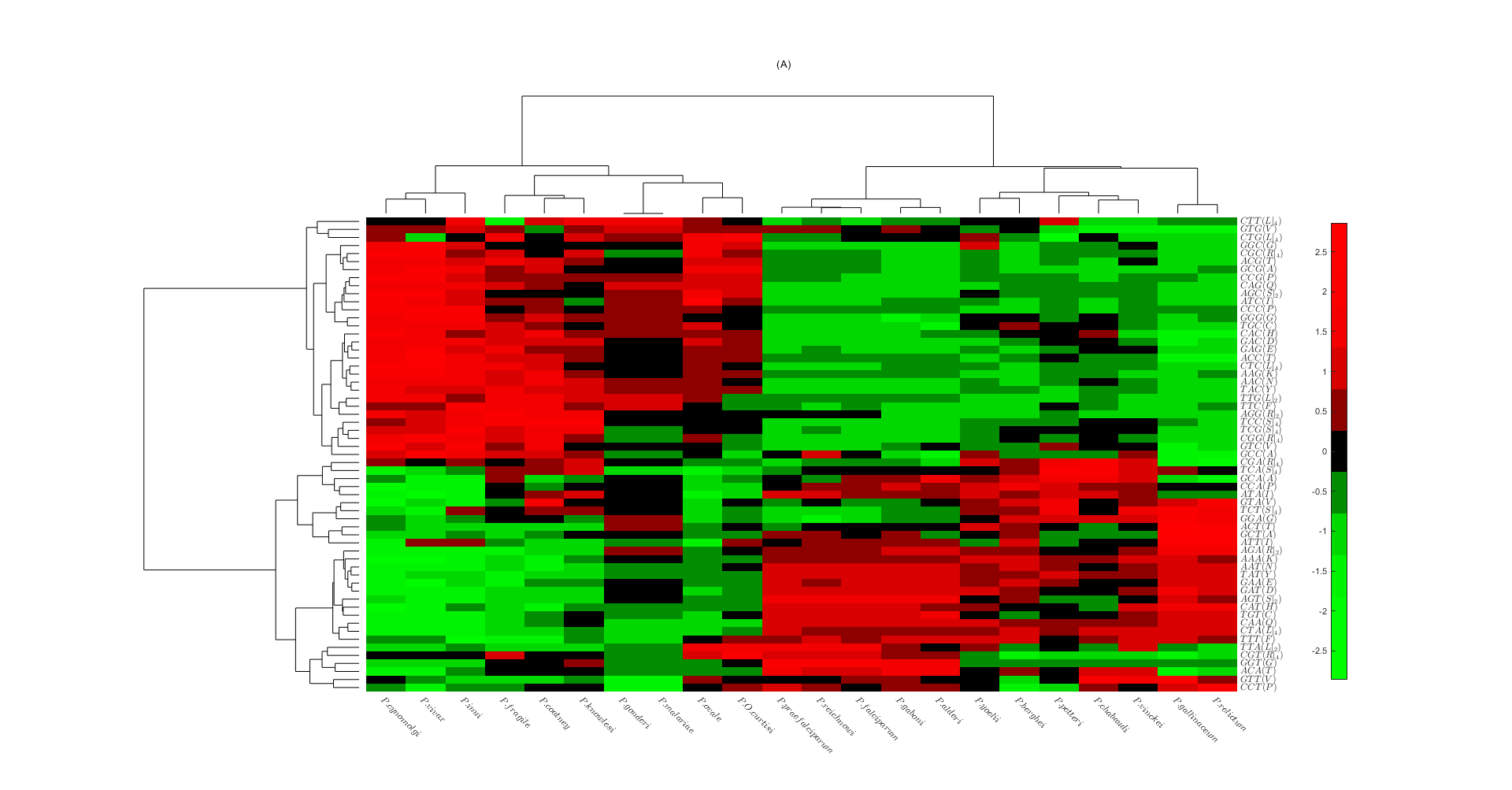

**Length and SPI results from other strain**

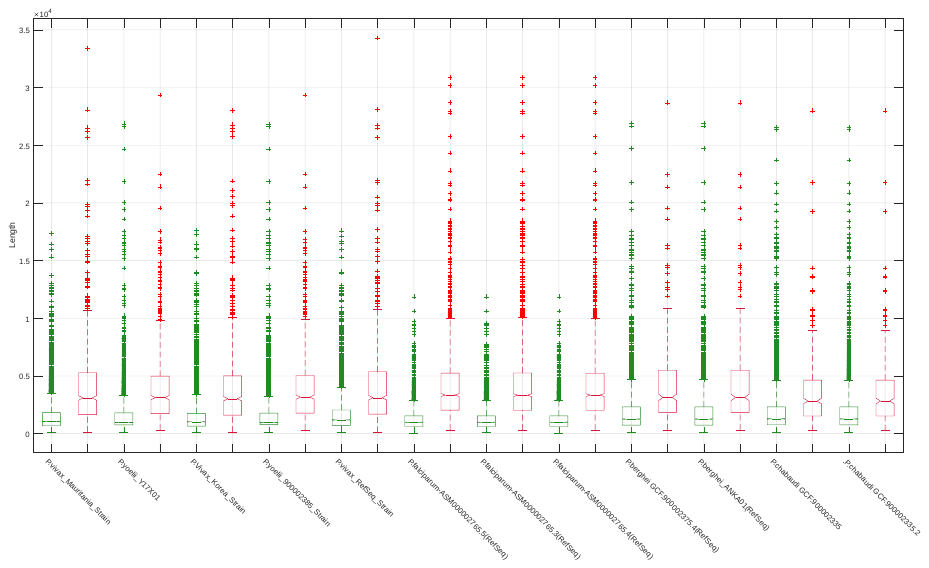

Fig. SM9 **LCPs are represented in red. nLCPs are represented in green. Length is measured in nucleotides.**

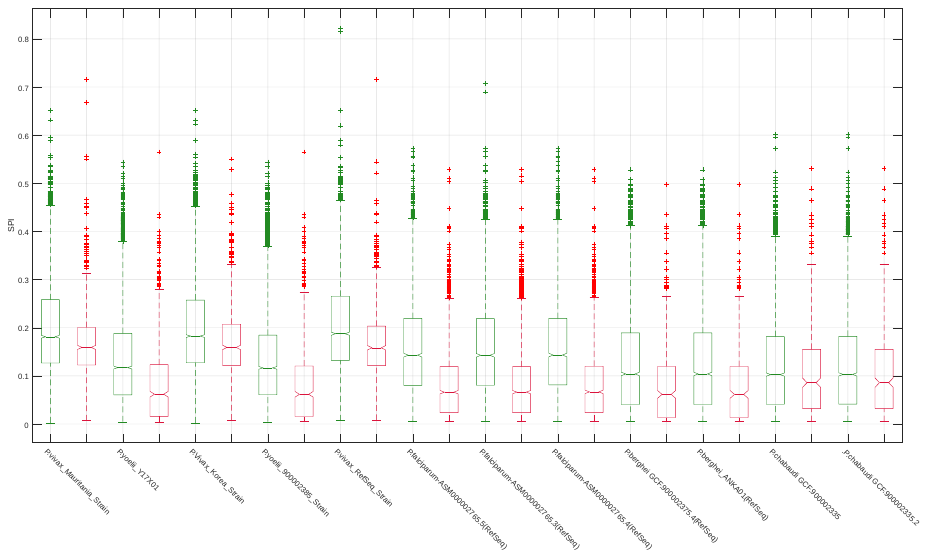

**Fig. SM10** LCPs are represented in red. nLCPs are represented in green

Herein, we considered different strains, when available, for some of the parasites analysed in the main corpus of the work. The split between LCPs and nLCPs returns also for these sets of genes both in length and SPI distributions.

**Length and Number of LCRs**

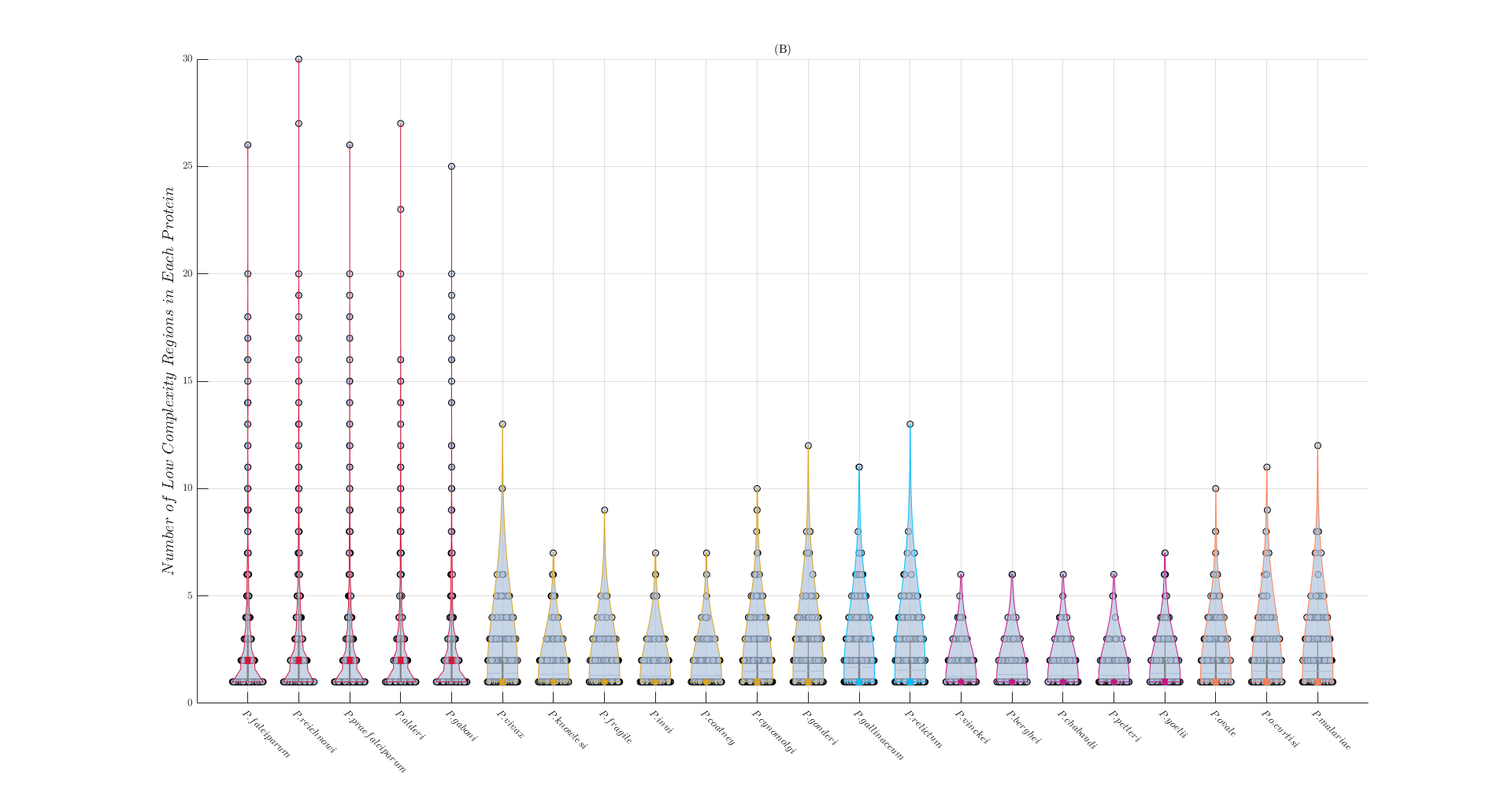

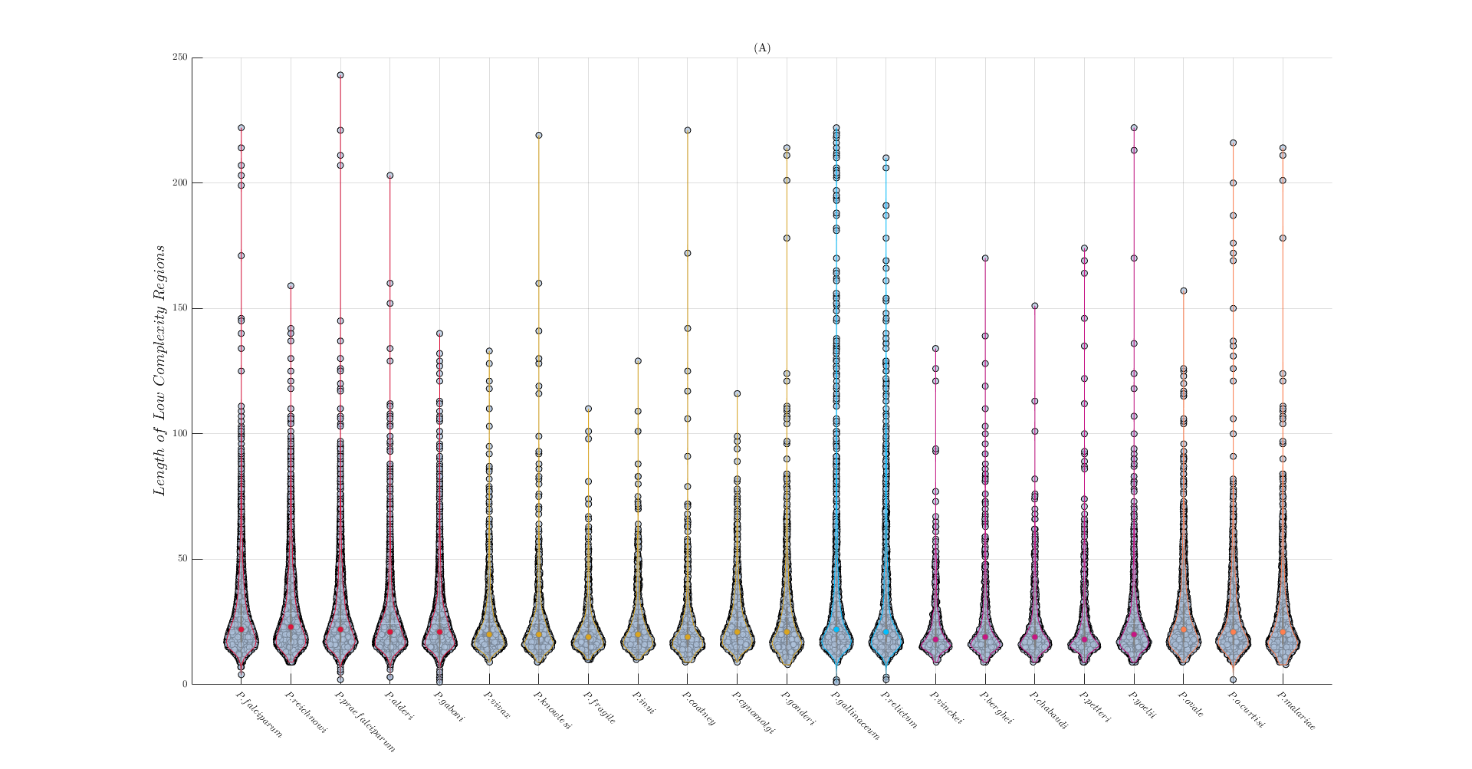

**Fig. SM11** (A) Length Distribution of LCRs (B) Number of LCRs in each protein

**Length and Number of Low Complexity Regions**

Length distributions of LCRs are represented by the violin plots in **Fig SM11(A)**. We performed MT2. LCRs of *P.falciparum, P.reichnowi* and *P.praefalciparum* are longer with significance from the rest of parasites (*p < 0.01*) made exception for *P.cynomolgi, P.gallinaceum, P.relictum, P.ovale* and *P.o.curtisi.* The number of LCRs of each *Plasmodium* are collected in **Tab.SM2.** We compared the distribution of each group (MT2). *Laverania Plasmodia* differ significantly from *Simian Plasmodia* and *Vinckeia* parasites. We do not find significant difference with *Haemamoeba* *Plasmodia* and HIPs. These last two comparisons are provided with the caveat that the statistical pauperism of these two groups could introduce a confound. Finally, we wondered how many LCRs sculp protein sequences on average (**Fig. SM11(B)**). *Laverania Plasmodia* incorporate the highest number of LCRs into their proteins, reaching, in some cases, 30 regions per protein. We have applied MT2. As expected, the distributions of *Laverania Plasmod*ia differ significantly from those of other parasites. Overall, *P.falciparum’s* family appears to be more likely to incorporate Low Complexity Regions into their Proteome than other parasites.

| **Tab. SM2** Number of LCRs in each plasmodium | |
| --- | --- |
| ***_Organism_*** | _Number of LCRs_ |
| ***_P.falciparum_*** | _4714_ |
| ***_P.reichnowi_*** | _4690_ |
| ***_P.praefalciparum_*** | _4637_ |
| ***_P.alderi_*** | _4337_ |
| ***_P.gaboni_*** | _4213_ |
| ***_P.vivax_*** | _1188_ |
| ***_P.knowlesi_*** | _795_ |
| ***_P.fragile_*** | _869_ |
| ***_P.inui_*** | _655_ |
| ***_P.coatney_*** | _658_ |
| ***_P.cynomolgi_*** | _1040_ |
| ***_P.gonderi_*** | _1176_ |
| ***_P.gallinaceum_*** | _1468_ |
| ***_P.relictum_*** | _1223_ |
| ***_P.vinckei_*** | _493_ |
| ***_P.berghei_*** | _564_ |
| ***_P.chabaudi_*** | _532_ |
| ***_P.petteri_*** | _500_ |
| ***_P.yoelii_*** | _1221_ |
| ***_P.malariae_*** | _1176_ |
| ***_P.ovale_*** | _1574_ |
| ***_P.o.curtisi_*** | _1161_ |

**Ratio between LCPs’ length and LCRs’ extension in different strains**

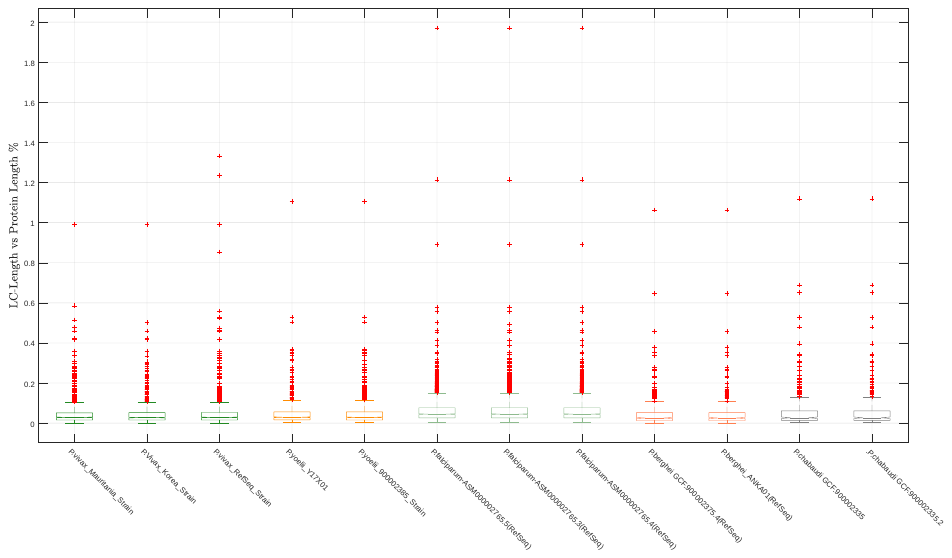

***Fig. SM12*** Proportion between length of LCRs and protein length in different Plasmodia strains

A feature that stands out from the observation of the boxplots is that compared to what was observed during the main dissertation of the work some of the parasites have different percentages of the ratio between the length of the proteins and the linear extension of the LCRs.

**Codon Contribution to LCRs’ Extension**

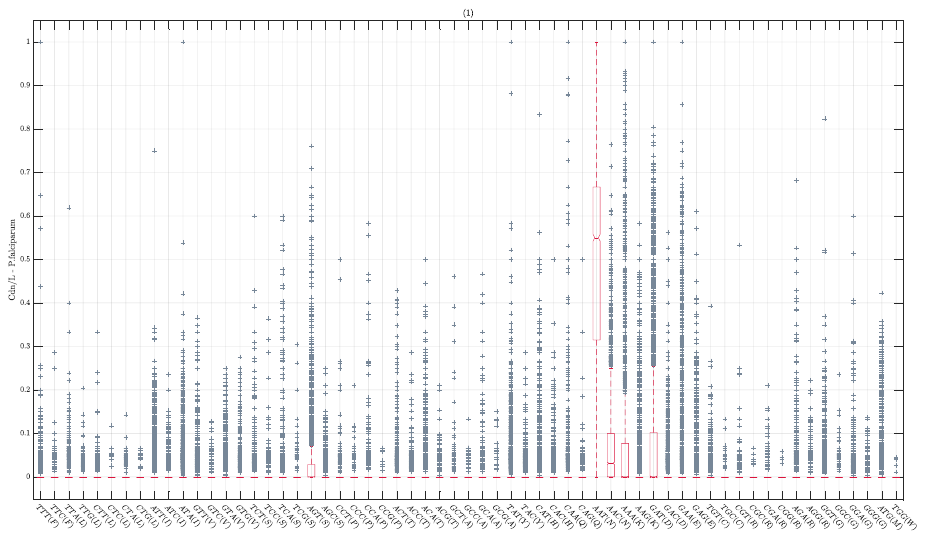

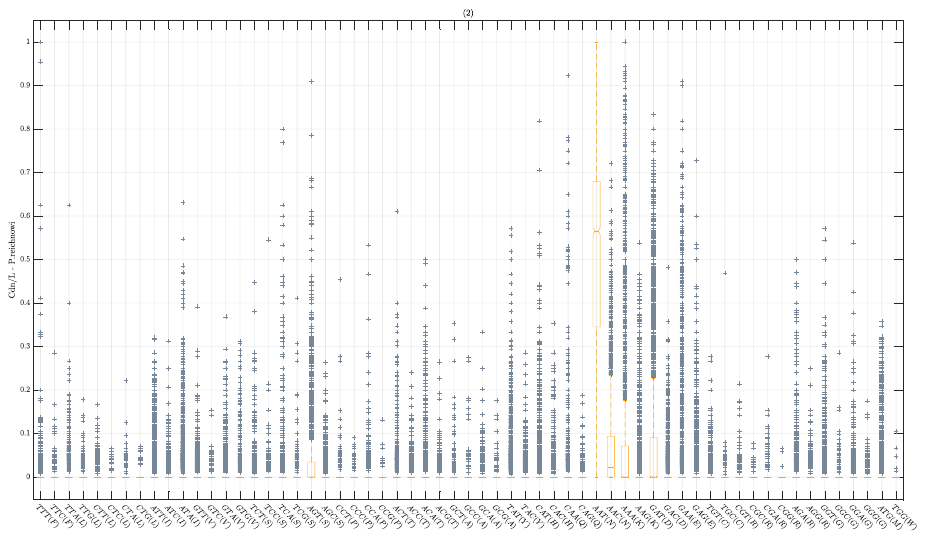

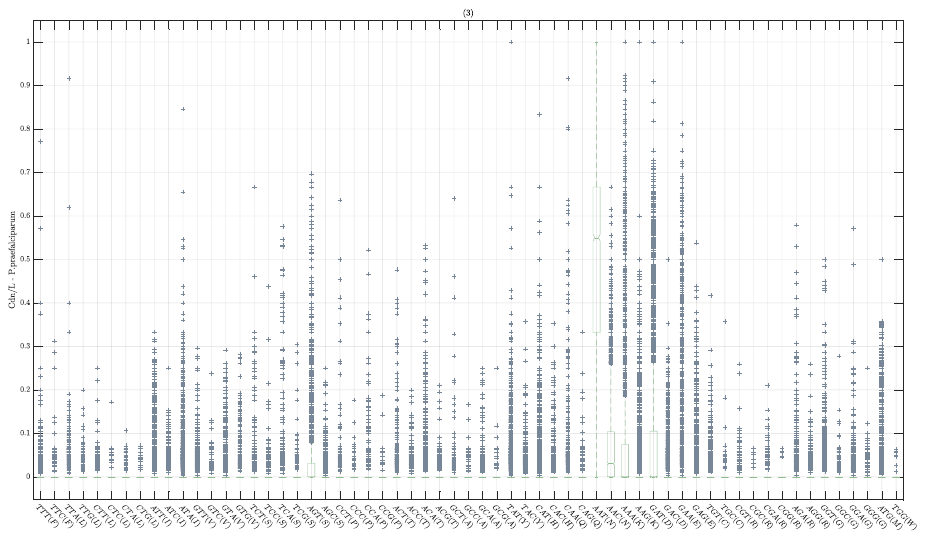

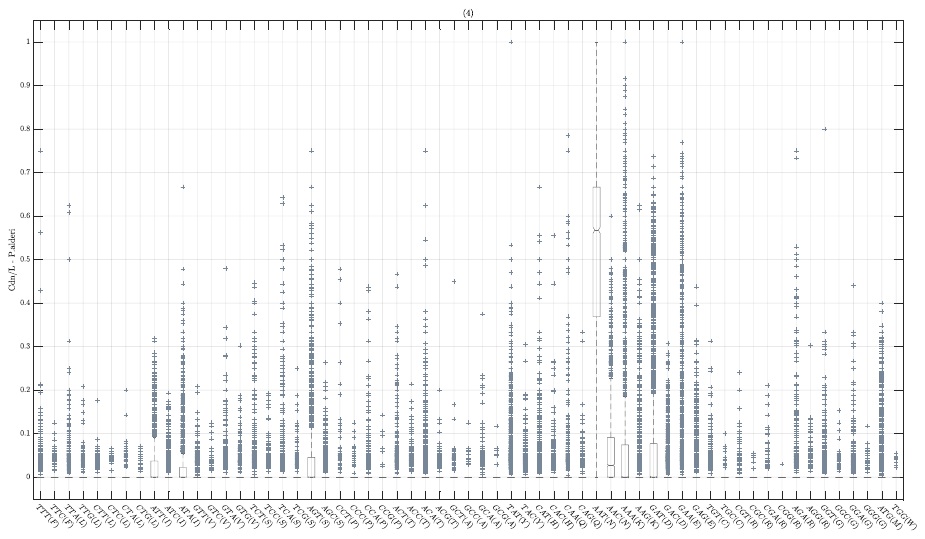

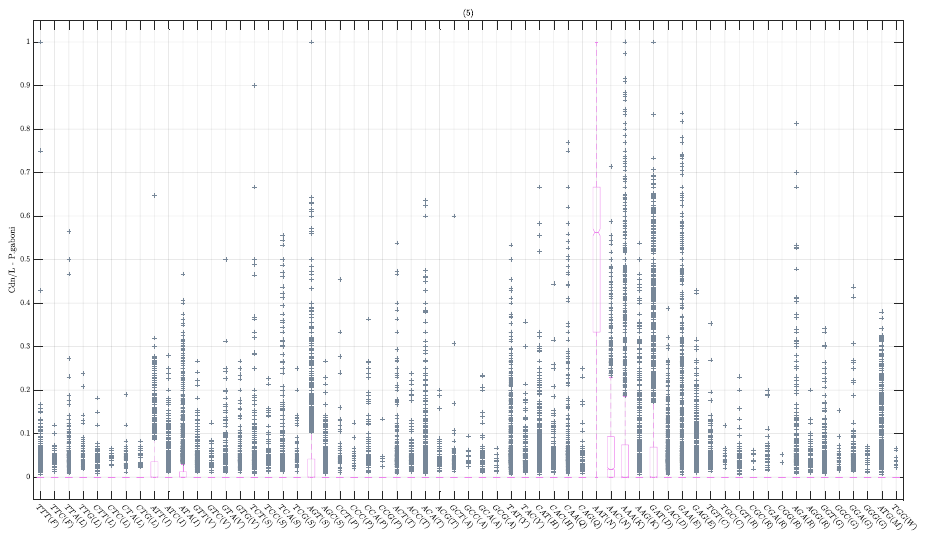

***Fig. SM13*** The image represents the composition of the LCRs of the Laverania plasmodia. We calculated the relationship between the quantity of a certain codon in a Low Complexity Region and the length of that region (relation hereafter referred as Cdn/L) . Unlike the other parasites, the Low Complexity regions represented here are mainly composed of AAT.

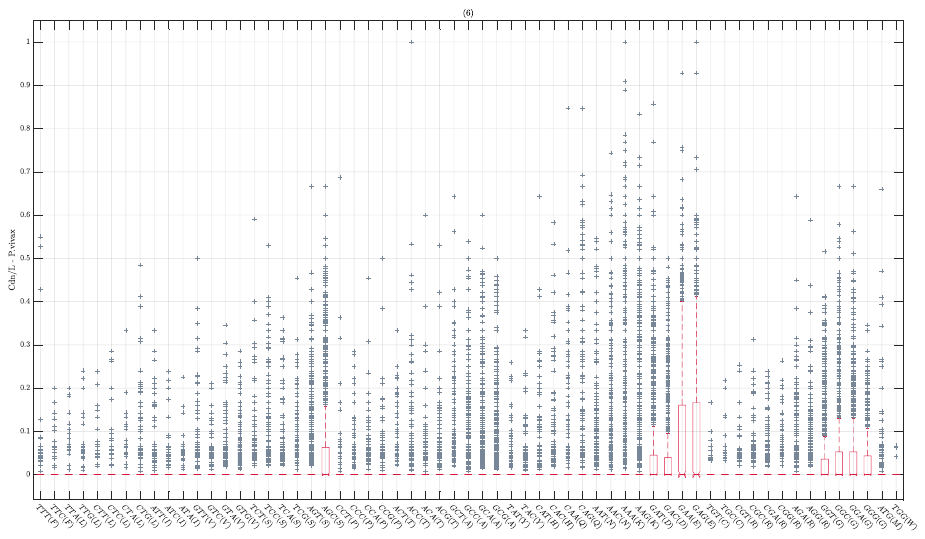

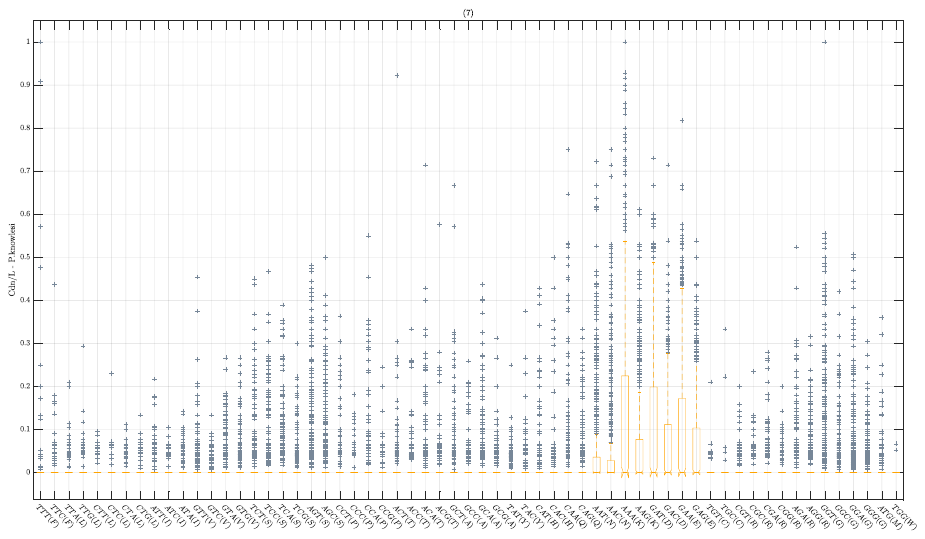

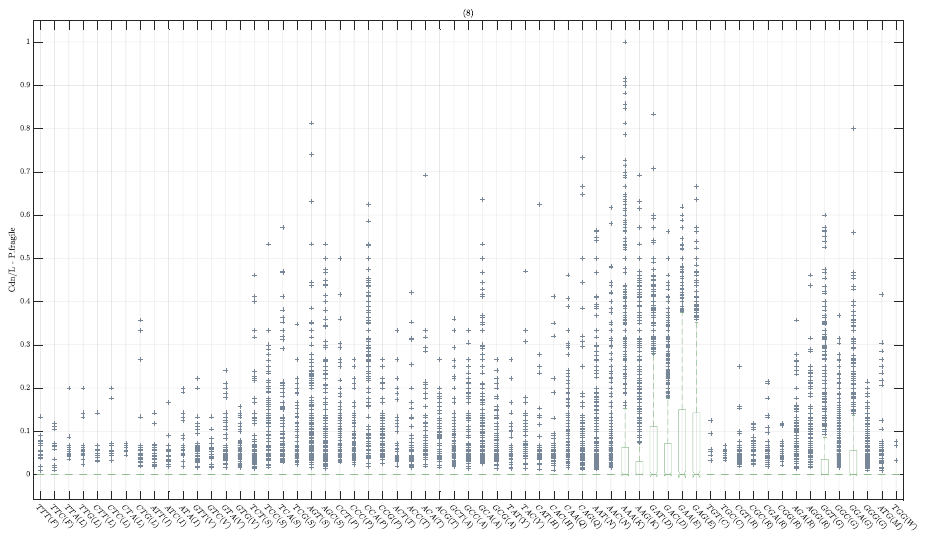

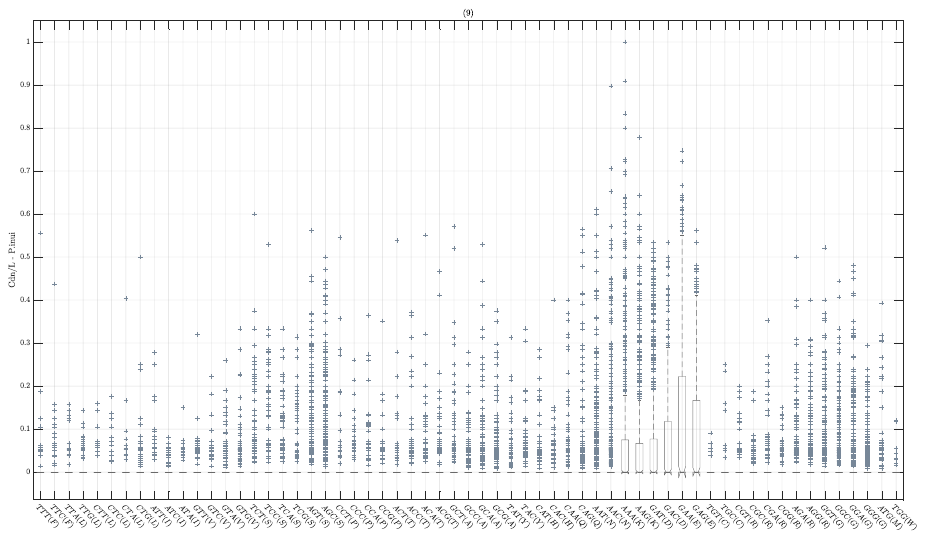

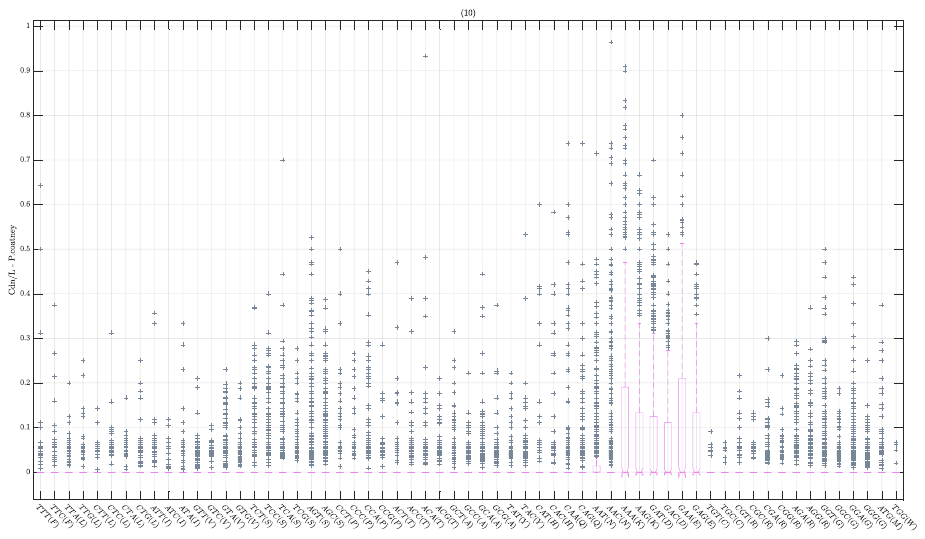

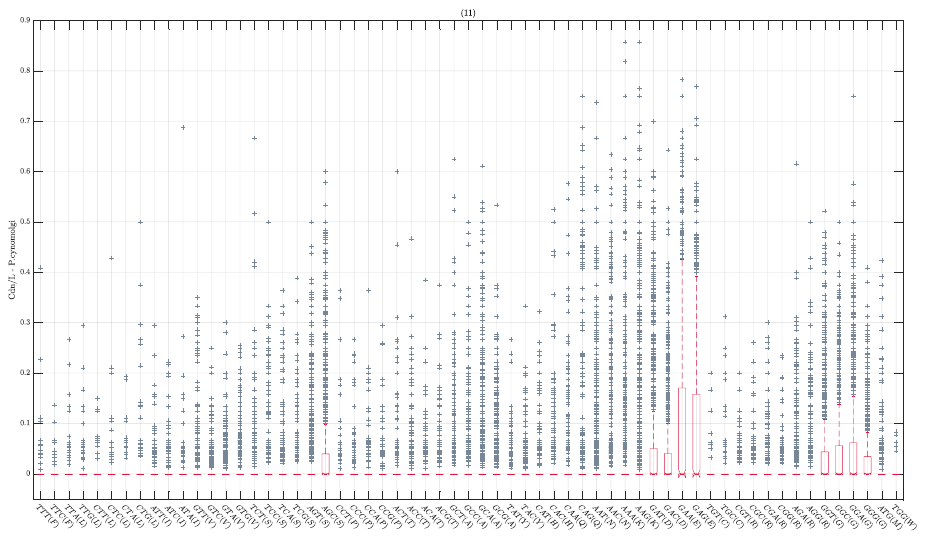

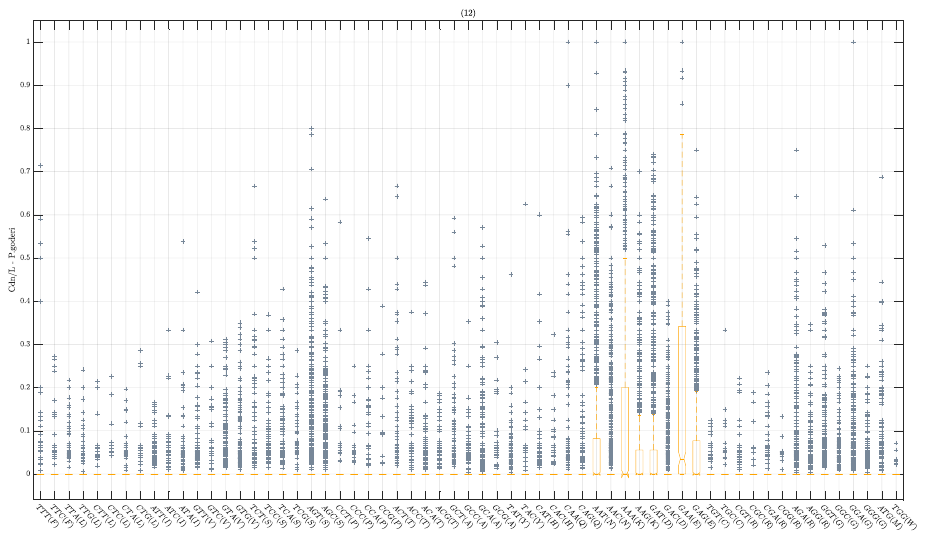

***Fig. SM14*** Cdn/L Simian Plasmodia

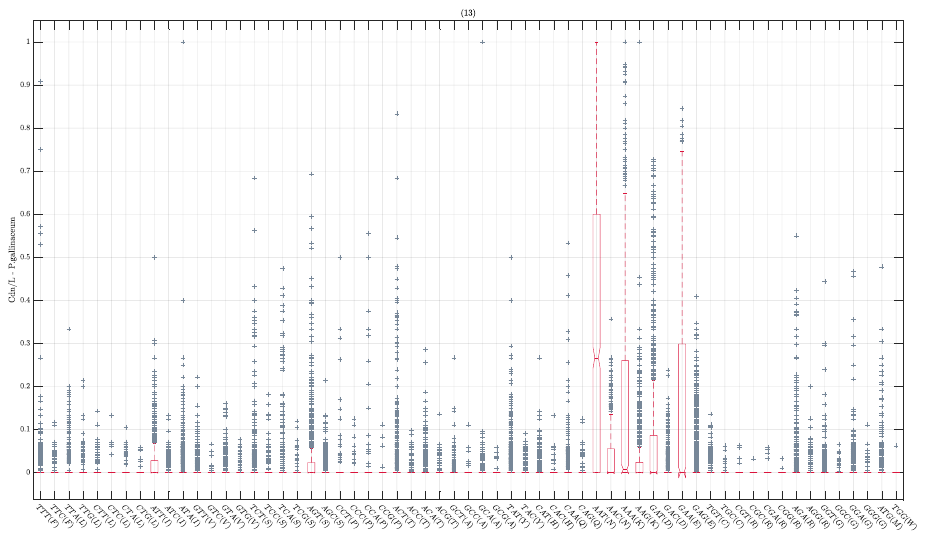

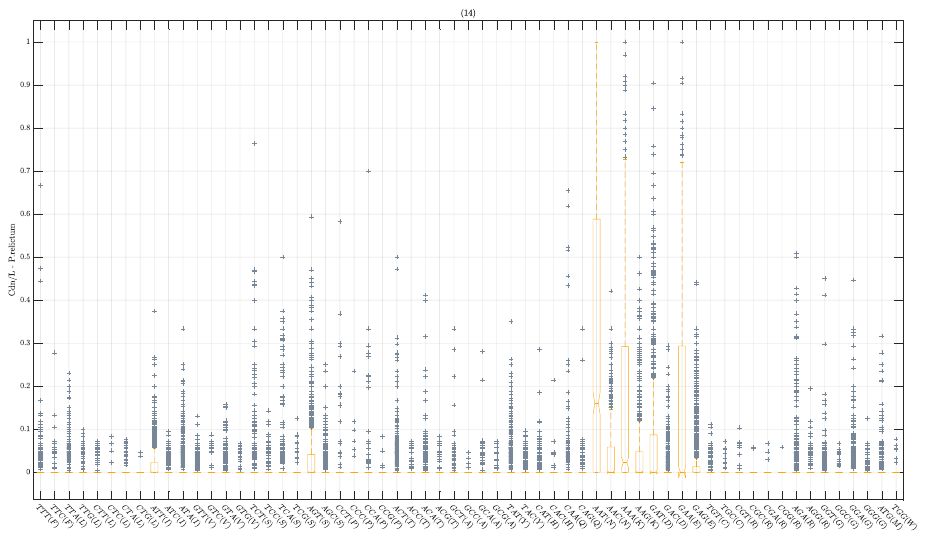

***Fig. SM15*** Haemamoeba Plasmodia

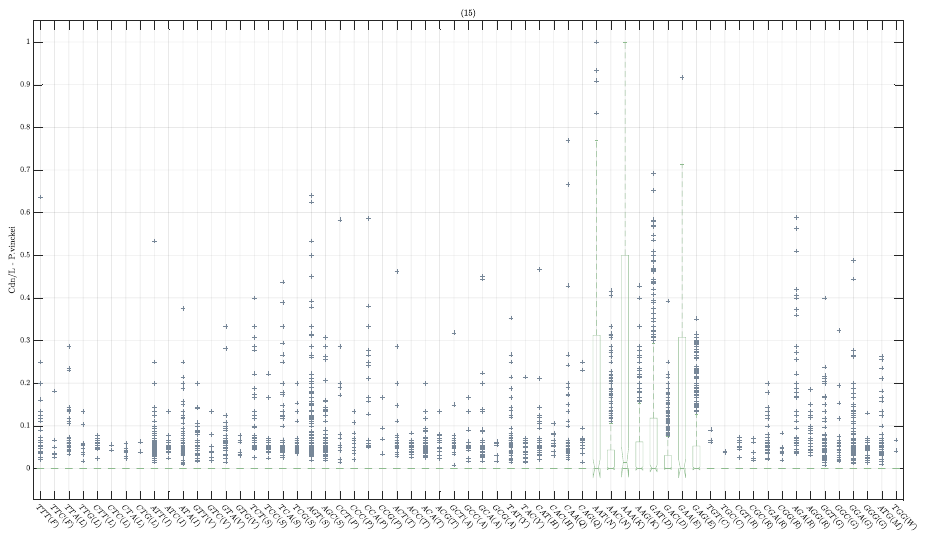

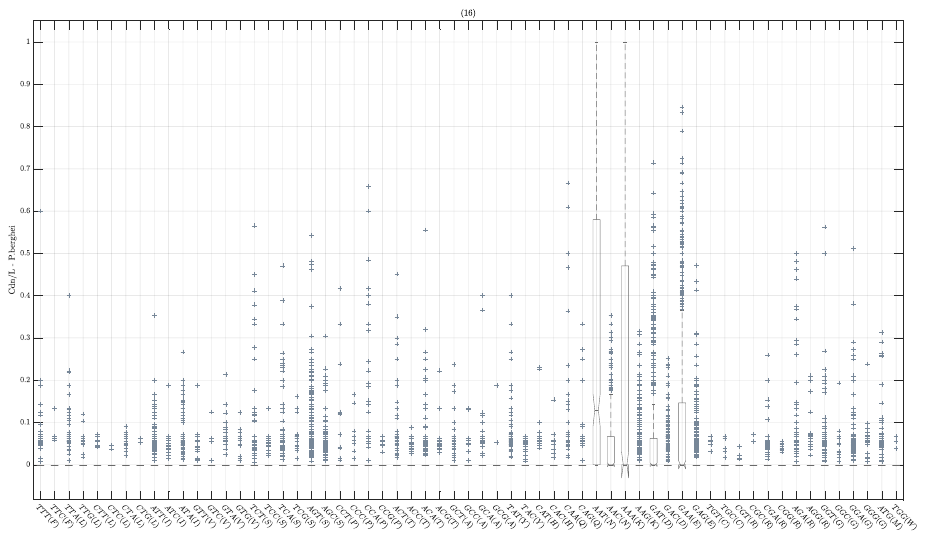

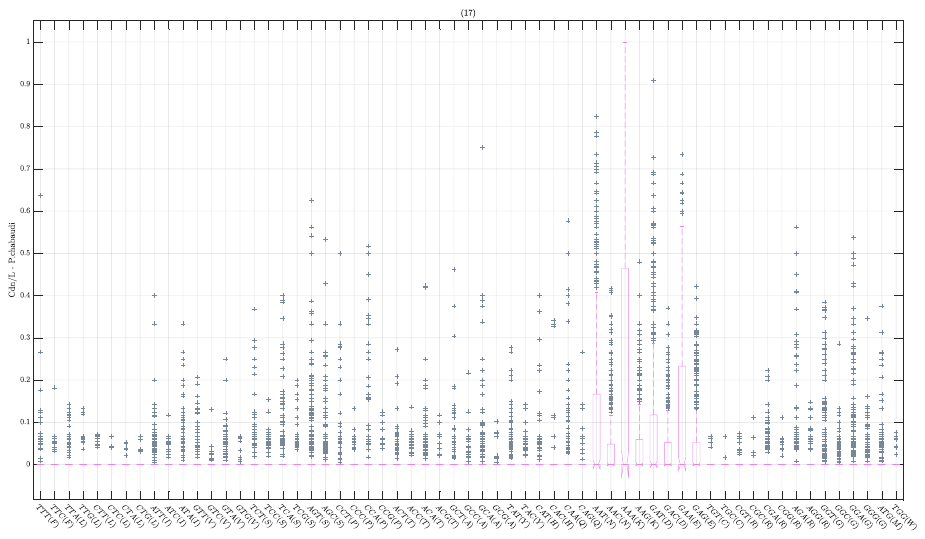

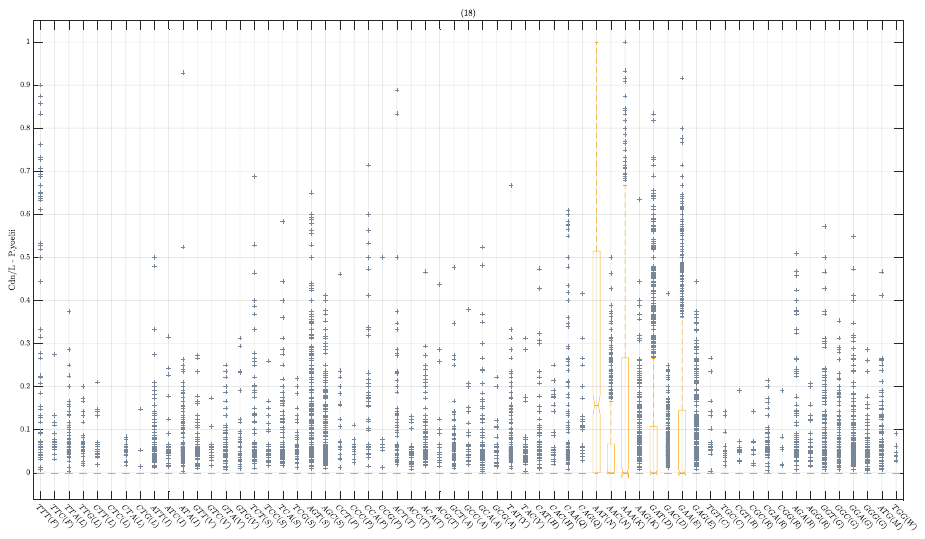

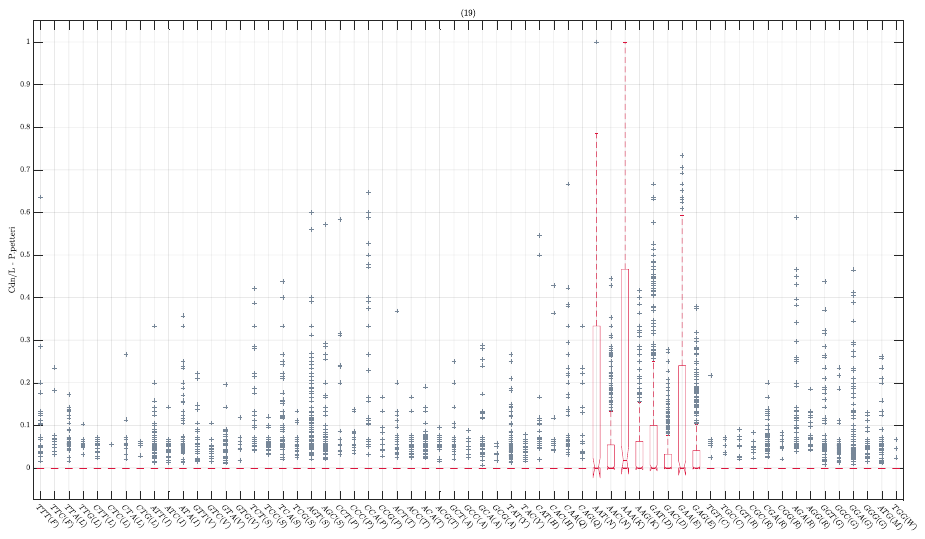

***Fig. SM16*** Vinckeia Plasmodia

***Fig. SM17*** HIPs

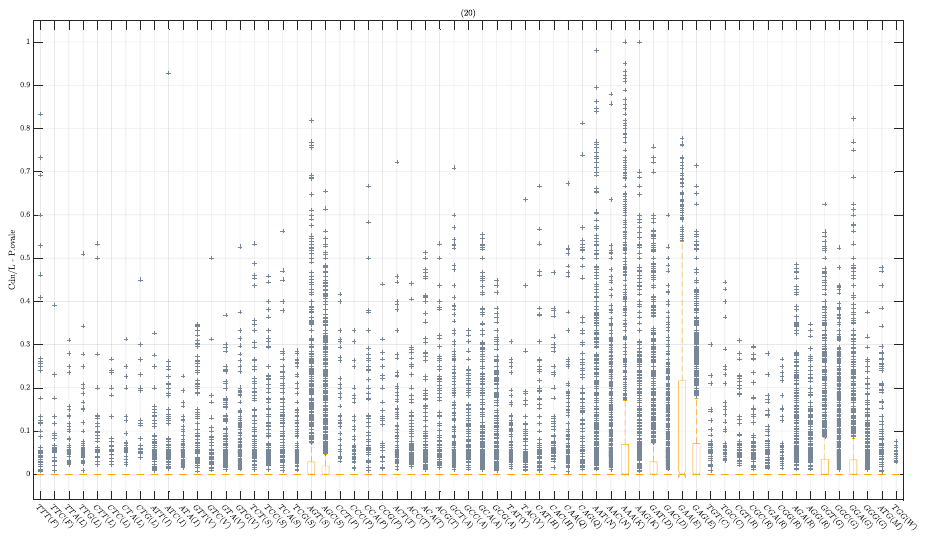

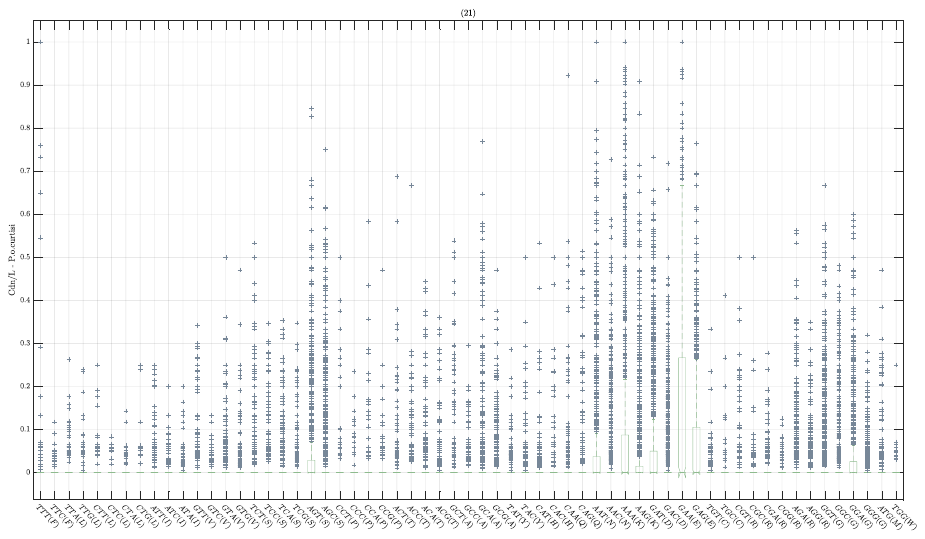

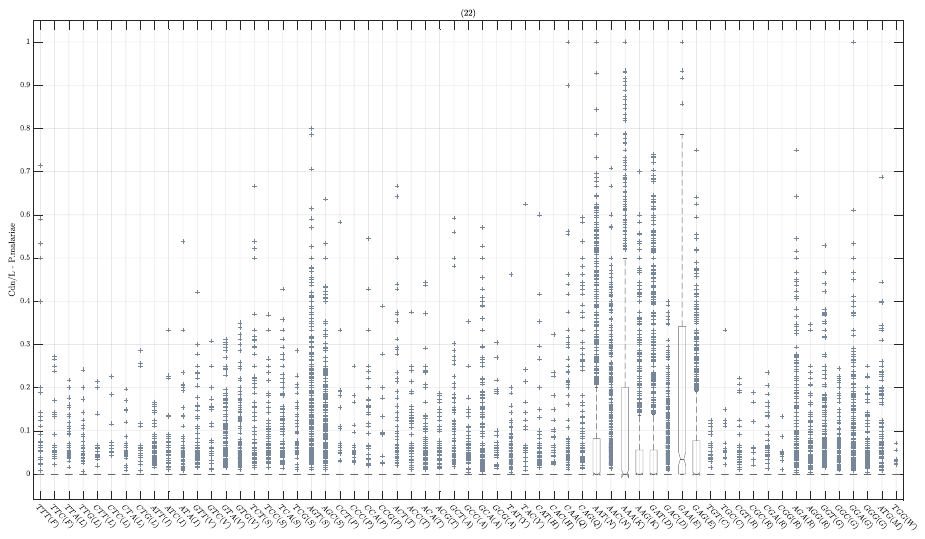

Each parasite presents a particular architecture for its Low Complexity Regions. Taking the same argument that we used for the calculation of the Shannon Entropy we have represented how much the same codon is used along the a low complexity region. Hence, if a point in the boxplot has value one, the LCR will be entirely composed of that codon.
